# *CHD8* Suppression Impacts on Histone H3 Lysine 36 Trimethylation and Alters RNA Alternative Splicing

**DOI:** 10.1101/2020.03.14.992032

**Authors:** Emanuela Kerschbamer, Takshashila Tripathi, Serkan Erdin, Elisa Salviato, Francesca Di Leva, Endre Sebestyén, Michele Arnoldi, Matteo Benelli, James F. Gusella, Silvano Piazza, Francesca Demichelis, Michael E. Talkowski, Francesco Ferrari, Marta Biagioli

## Abstract

Disruptive mutations in the chromodomain helicase DNA binding protein 8 (*CHD8*) have been recurrently associated with Autism Spectrum Disorders (ASD). In normal cellular physiology, CHD8 co-purifies with MLL1 and MOF transcriptional activation complex, with elongating RNAPII and directly binds to DNA promoters and enhancers regions, thus a regulatory role in transcriptional initiation and elongation could be postulated.

Here we investigated how chromatin landscape reacts to *CHD8* suppression by analyzing a panel of histone modifications in induced pluripotent stem cell-derived neural progenitors. We interrogated transcriptionally active and repressed regions, as well as active and poised enhancers.

*CHD8* suppression led to significant reduction (47.82%) in histone H3K36me3 peaks at gene bodies, particularly impacting on transcriptional elongation chromatin states. H3K36me3 reduction specifically affects highly expressed, CHD8-bound genes. Histone H3K36me3 reduction associated to *CHD8*-suppression does not functionally impact on global transcriptional levels, but correlated with altered alternative splicing patterns of ∼ 2000 protein coding genes implicated in “RNA splicing”, “mitotic cell cycle phase transition” and “mRNA processing”, especially affecting alternative first exon and exon skipping events.

In summary, our results point toward broad molecular consequences of *CHD8* suppression, implicating altered histone deposition/maintenance and RNA processing regulation as important regulatory processes in ASD.

## Introduction

*De novo* truncating mutations in *CHD8* have been reported and independently validated to be a strong risk factor for autism spectrum disorders (ASD) (Neale et al. 2012; O’Roak et al. 2012a; O’Roak et al. 2012b; Talkowski et al. 2012; Iossifov et al. 2014; Parikshak et al. 2013; De Rubeis et al. 2014). *CHD8* has been classified as a high confidence ASD candidate risk factor (score 1) in the Simons Foundation Autism Research Initiative (SFARI) [https://gene.sfari.org (Abrahams et al. 2013)] with most *CHD8* mutations, identified from ∼70 ASD probands (www.ncbi.nlm.nih.gov/clinvar/ interrogated on January 8^th^, 2020), considered disruptive and resulting in protein haploinsufficiency. *CHD8* defines a subclass of ASD patients, displaying evident macrocephaly, distinct faces, sleep problems and gastrointestinal complaints (Bernier et al. 2014; Yasin et al. 2019). Most of these phenotypic characteristics were recapitulated in *chd8* knock-down zebrafish (Bernier et al. 2014; Sugathan et al. 2014) and, more recently, in *Chd8* suppression mouse models (Durak et al. 2016; Katayama et al. 2016; Suetterlin et al. 2018). Indeed, *chd8*-morpholino zebrafish and *Chd8* heterozygous mice display increased brain size, due to increased numbers of proliferating cells and newborn neurons, possibly initiated by altered gene expression in the developing neocortex (Sugathan et al. 2014; Suetterlin et al. 2018). Remarkably, genome-wide transcriptomic changes that impact on ASD related genes were also detected *in vitro*, in human neural progenitor cells (NPC) with reduced *CHD8* expression (Sugathan et al. 2014). Taken together, these observations suggest that aberrant genome-wide transcription leading to altered brain development is strictly correlated to reduced levels of CHD8 function. However, the detailed molecular mechanism through which CHD8 regulates this process still remains obscure.

A direct effect can be proposed since CHD8 is able to bind DNA at promoters and enhancer regions in hNPCs, mouse midfetal brain and embryonic cortex (Sugathan et al. 2014; Cotney et al. 2015). However, an indirect mechanism can also be postulated since other genes, not bound by CHD8, appear also to be transcriptionally dysregulated following *CHD8* suppression (Sugathan et al. 2014). Chromatin structure is intimately related to transcription, as DNA tightly packed around nucleosomes prevents transcription, while, conversely, the exchange or removal of nucleosomes allows free access to DNA, thus correlating with active gene expression (Venkatesh and Workman 2015). Using mass spectrometry, CHD8 was shown to co-purify with components of the MLL and CoREST, SWI/SNF and NuRD ATP-dependent remodeling complexes, supporting its possible role in transcriptional initiation (Thompson et al. 2008). On the other hand, CHD1, another member of the CHD protein family, has been shown to interact with the PAF1 transcription elongation complex maintaining H3K4me3/H3K36me3 domains at actively transcribed genes (Lee et al. 2017). Reduction of CHD1 alters H3K4me3 and H3K36me3 patterns, suggesting a role for CHD1 in establishing/maintaining the boundaries of these mutually exclusive histone marks (Lee et al. 2017). The SETD2 and SETD5 histone H3, Lysine 36 methyltransferases normally associate with RNAPII, and their activity results in increased H3K36me3 toward the 3’ end of active genes (Krogan et al. 2003; Kizer et al. 2005). Loss of SETD2/5-dependent H3K36me in mammals (Sessa et al. 2019) as well as loss of yeast Chd1 causes reduced Rpd3S activity (histone deacetylase complex), increased acetylation, and increased cryptic transcription within gene bodies (Selth et al. 2010). Based on its placement on the phylogenetic tree and the presence of an ATPase domain (Thompson et al. 2008), CHD8 is most likely acting as an ATP-dependent chromatin remodeling factor; thus, similar to CHD1, CHD8 loss might cause increased nucleosome turnover and alterations in co-transcriptional processes, such as cryptic transcription within gene bodies and alternative splicing (Radman-Livaja et al. 2012; Smolle et al. 2012).

Alternative splicing (AS), one of the major contributor to protein diversity in eukaryotes, is important for highly specialized cells, such as neurons. Aberrant splicing might contribute to neuronal dysfunction and has been associated with several neurological diseases (Cuajungco et al. 2003; Mordes et al. 2006) (Wang et al. 2012; Gompers et al. 2017). Broad chromatin conformation and transcriptional kinetics are major factors in the regulation of AS. Chromatin relaxation accelerates RNAPII processing and correlates with alternative exons skipping; conversely, packed nucleosomes slow down RNAPII progression causing pausing of transcription and the inclusion of non-constitutive weak exons (Luco et al. 2010; Nilsen and Graveley 2010; Luco and Misteli 2011). As experimental evidence is pointing to dysregulated chromatin regulation as a key feature in the pathogenesis of ASD, it is tempting to hypothesize that chromatin function of CHD proteins, and CHD8 in particular, might act to regulate RNA transcription, elongation and processing thereby being responsible for the characteristic neurodevelopmental effects observed in ASD.

In order to possibly dissect this mechanism, we characterized the consequences of *CHD8* suppression on the chromatin landscape, analyzing different histone modifications using chromatin immunoprecipitation and sequencing (ChIP-seq). Specifically, we interrogated histone marks characteristic of transcriptionally active (H3K4me2, H3K4me3, H3K27ac, H3K36me3) and repressed regions (H3K27me3) as well as active enhancers (H3K4me1, H3K27ac) in control induced pluripotent stem cell (iPSC)-derived neuronal progenitors cell line and in previously characterized lines where a ∼50% reduction in *CHD8* was obtained by lentiviral delivery of shRNAs (Sugathan et al. 2014). We uncovered alterations affecting the H3K36me3 histone mark in the body of highly transcribed genes, which do not primarily affect RNA transcription, but rather alter alternative splicing of genes implicated in “histone modification”, “covalent chromatin modification” and “mitotic cell cycle phase transition”, thus unveiling altered histone deposition/maintenance and RNA processing regulation as important regulatory processes in ASD.

## Results

### Epigenetic consequences of CHD8 suppression in human neuronal progenitor cells

To assess the functional consequences of *CHD8* suppression in chromatin organization, we resourced to previously characterized control iPSC-derived neuronal progenitors cell line, GM8330-8 and its derivatives where ∼50% reduction in *CHD8* was obtained by lentiviral-mediated delivery of short-hairpin RNAs (shRNAs) (Sugathan et al. 2014). In these model systems, we analyzed six different histone modifications using chromatin immunoprecipitation and sequencing (ChIP-seq), specifically interrogating transcriptionally active (H3K4me2, H3K4me3, H3K27ac, H3K36me3) and repressed regions (H3K27me3) as well as active enhancers (H3K4me1, H3K27ac). For each of the six histone marks, three independent shRNAs targeting the coding sequence of *CHD8* (Sh1, Sh2, Sh4) and two technical replicate controls against *GFP* sequence were used (Fig. 1A). Sh1-*CHD8*, Sh2-*CHD8*, Sh4-*CHD8* presented nearly comparable levels of CHD8 at ∼ 50% of the physiological levels, thus precisely mimicking the human haploinsufficiency condition (Sugathan et al. 2014). Importantly, for *CHD8*-knock-down models as well as for *GFP* controls, genome-wide transcriptomic data and CHD8 binding sites were available (Fig. 1A) (Sugathan et al. 2014).

**Figure 1.**
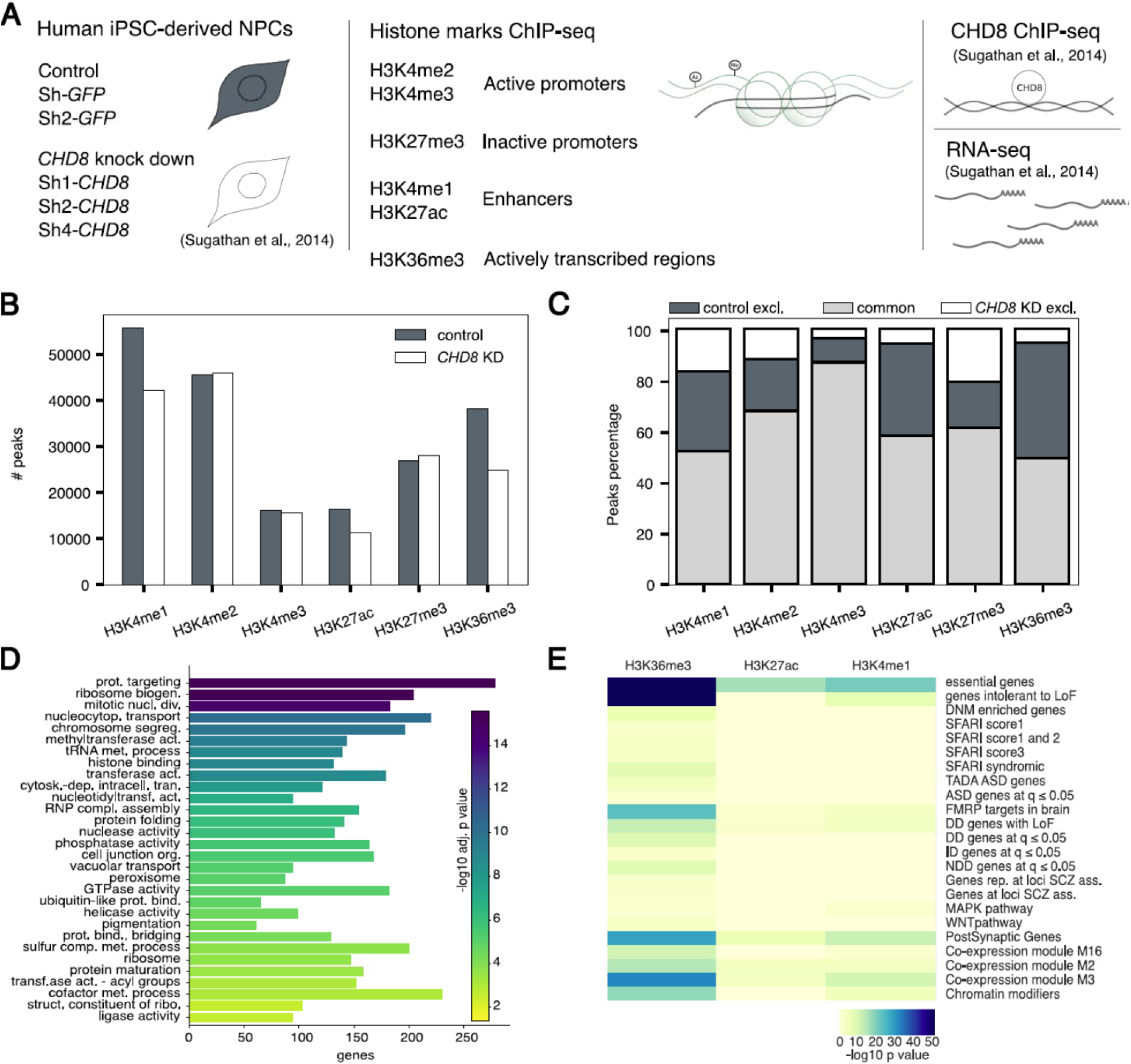
Altered chromatin landscape upon CHD8-suppression: reduction of histone H3K27ac, H3K36me3 and H3K4me1. A. Schematic representation of the study design and integrative approach used in this work. Human iPSC-derived NPCs (hiNPC) knocked-down for *CHD8* (Sh1-, Sh2- and Sh4-*CHD8*) and control hiNPCs (Sh-*GFP,* Sh-*GFP2*) (Sugathan et al. 2014), were analyzed via ChIP-seq for six histone marks representative of different chromatin regions - active promoters (H3K4me2, H3K4me3), inactive promoters (H3K27me3), enhancers (H3K4me1, H3K27ac) and actively transcribed regions (H3K36me3). ChIP-seq results were subsequently integrated with CHD8 binding sites and available transcriptomics (RNA-seq) datasets obtained from the same model system (Sugathan et al. 2014). B. The bar plot reports the total number of peaks identified in controls (grey) and *CHD8* knock-downs (white) for each histone mark analyzed. The y-axis presents the number of peaks resulting from the intersection of two replicates for each experimental condition: controls (intersection between Sh-*GFP,* Sh-*GFP2*), CHD8-knock-down (intersection between Sh2 and Sh4 *CHD8*). Histone marks H3K4me1, H3K27ac and H3K36me3 were the mostly affected by *CHD8* knock-down, displaying, in all cases, decreased number of peaks in *CHD8* suppression hiNPC models (19388, 6445 and 20507 peaks were lost, respectively). C. The bar plot highlights the percentage of peaks in common between controls (intersection between Sh-*GFP,* Sh-*GFP2*) and CHD8-knock-down (intersection between Sh2- and Sh4-*CHD8*) (light gray). Percentage of peaks exclusively present in controls (intersection between Sh-*GFP,* Sh-*GFP2*) or in *CHD8* knock-down (intersection between Sh2- and Sh4-*CHD8*) are presented in grey and white, respectively. The total number of peaks (100%) is the sum of the peaks common and exclusive to each experimental condition. Upon *CHD8* suppression, H3K4me1, H3K27ac and H3K36me3 confirm extensive peak loss compared to controls, with a reduction of 1.22%, 36.23% and 45.20%, respectively. D. Graphical representation of slim GO functional annotation enrichment for all genes interested by H3K4me1, H3K27ac and H3K36me3 peaks depletion following *CHD8* knock-down. Top 30 GO terms are shown in figure, while full list of significant terms is reported in Suppl. Table 1. Color-coded bar plot according to adjusted p values in -log10 scale, displays statistically significant slim GOs. The size of the bar (x-axis) reports the number of genes for each slim GO term. E. The heatmap represents gene set enrichment p values in -log10 scale for all genes losing H3K4me1, H3K27ac and H3K36me3 peaks following *CHD8* knock-down. Genes’ lists related to ASD, neurodevelopment, co-expression modules in brain and intolerance to loss of function, were tested for enrichment as described in Materials and Methods. Full gene list description and enrichment results are available in Supplementary Table 1.

ChIP-seq experiments were conducted to obtain on average 40M reads for narrow marks (H3K4me3/me2/me1, H3K27ac) and 60M for broad histone marks (H3K36me3/K27me3) and for INPUT samples. After mapping and filtering, an average of 42308 peaks per sample was identified (Suppl. Fig. 1A-B), and showed the expected enrichment pattern and metagene profiles at the transcriptional start site (H3K4me3/me2/me1, H3K27ac), gene body (H3K36me3) of actively transcribed genes or on larger genomic regions spanning transcriptionally silent gene units (H3K27me3) and the expected enrichment correlation with ENCODE public datasets (Suppl. Fig. 1C-D) (Consortium 2012; Davis et al. 2018).

Upon *CHD8* suppression, H3K4me1, H3K27ac and H3K36me3 presented a substantial decrease in the number of peaks (37.44%, 38.45% and 47.82%, respectively) compared to control (intersection of Sh-*GFP* and Sh-*GFP2*) (Fig. 1B-C). Peaks detected exclusively in knock-down replicates (intersection of Sh2-*CHD8* and Sh4-*CHD8*, the two Sh-*CHD8* presenting nearly identical (46.63-48.61%) levels of *CHD8* suppression (Fig. 2C-D)) were limited (Fig. 1C). Notably, the third biological replicate Sh1-*CHD8*, presenting lower levels of *CHD8* suppression (Fig. 2C-D) (Sugathan et al. 2014), generally confirmed previous findings and the intersection of two biological replicates - in any of the possible combinations (Sh1-Sh2 *CHD8*, Sh2-Sh4 *CHD8*, Sh1-Sh4 *CHD8*), supported the reduction in H3K4me1, H3K27ac and H3K36me3 enrichment peaks (Suppl. Fig. 2A). In addition, Sh2-*CHD8* and Sh-*GFP* independent ChIP-seq datasets (generated in different laboratory than previous set) again sustained the conclusion that *CHD8* suppression was associated with a decreased H3K4me1 and H3K36me3 enrichment (Suppl. Fig. 2B).

**Figure 2.**
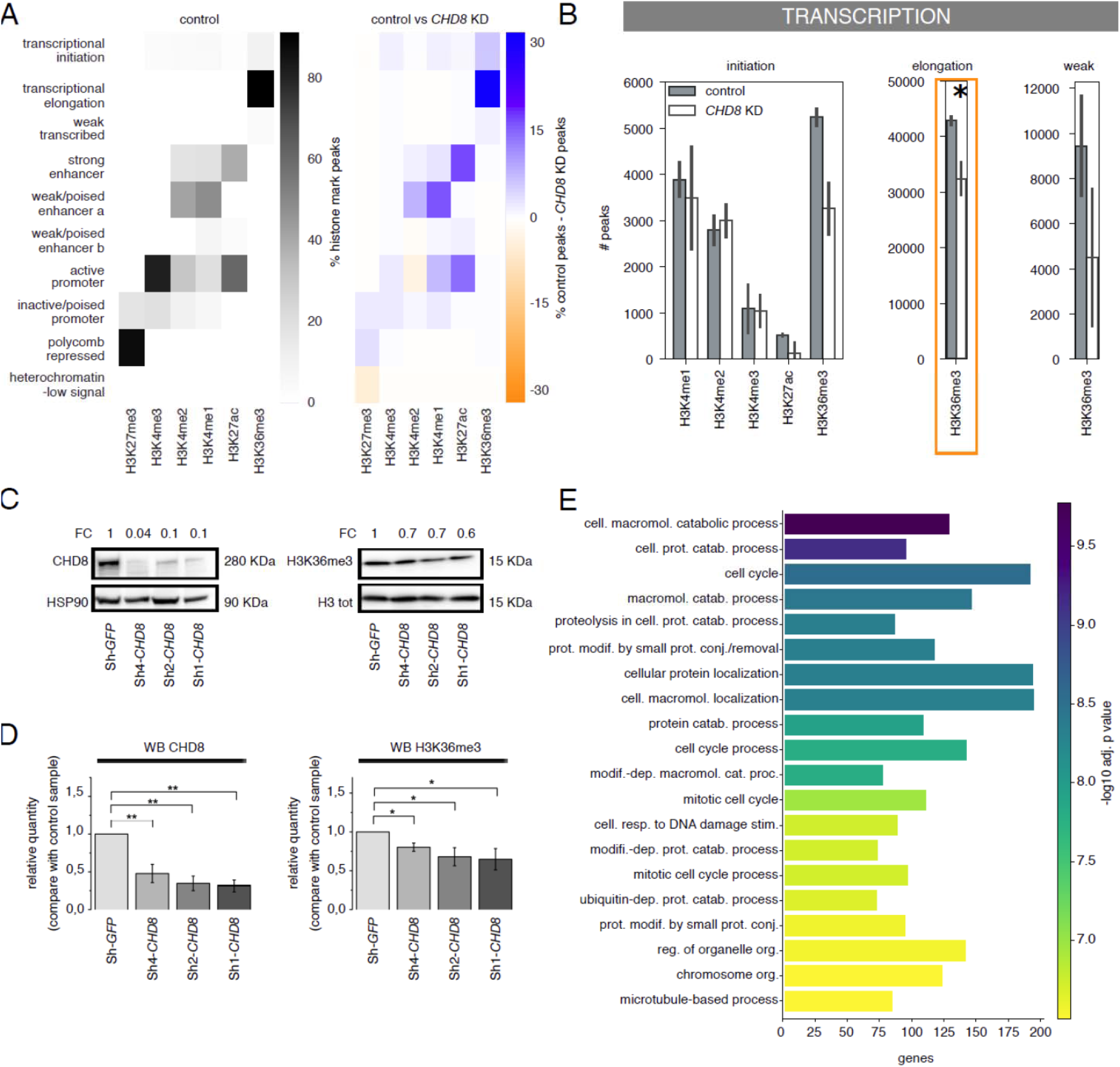
CHD8-suppression significantly impacts on histone H3K36me3 enrichment at transcriptional elongation sites. A. The heatmaps represent 10 different chromatin states (1. transcriptional initiation, 2. transcriptional elongation, 3. weakly transcribed, 4. strong enhancer, 5. weak/poised enhancer a, 6. weak/poised enhancer b, 7. active promoter, 8. inactive/poised promoter, 9. polycomb repressed, 10. heterochromatin/low signal), defined by the combination of different histone marks in control hiNPC as defined by ChromHMM(Ernst and Kellis 2012). The distribution of histone marks peaks across different chromatin states (see materials and methods for details) is presented as percentage of the total and color-coded in the heatmap (left). On the right, the heatmap describes the difference in number of peaks between two experimental conditions (controls versus *CHD8* knock-down). In blue are indicated chromatin states enriched in control, in orange chromatin states enriched in *CHD8* knock-down. H3K36me3 in transcriptional elongation is identified as the most affected chromatin state. B. The bar plots represent the number of peaks for each histone mark identified at transcriptional initiation (left), elongation (center) and weakly transcribed (right) genomic regions. Grey bars indicate controls (n=2, Sh-*GFP,* Sh-*GFP2*) and white bars refer to *CHD8* knock-down (n=2, Sh2-*CHD8*, Sh4-*CHD8*). H3K36me3 peak loss upon *CHD8* suppression was significant at the transcriptional elongation states (two biological replicates, T-test, p<0.05). C. The image describes a representative western blot illustrating CHD8 (left) and total histone H3K36me3 (right) levels across control (Sh-*GFP*) and *CHD8* knock-down clones (Sh1-*CHD8*, Sh2-*CHD8*, Sh4-*CHD8*). Down-regulation of CHD8 and total H3K36me3 amount is observed following *CHD8* suppression. Levels of CHD8 and H3K36me3 reduction are indicated in the top part of the panel as fold change (FC) compared to control Sh-GFP. Comparable amounts of total protein were loaded and HSP90 and total histone H3 was used as loading control. D. The bar chart reports fold change differences in CHD8 (left) and histone H3K36me3 left) levels comparing control (Sh-*GFP*) and *CHD8* knock-down clones (Sh1-*CHD8*, Sh2-*CHD8*, Sh4-*CHD8*) in western blotting experiments. The bars represent normalized (HSP90 and total histone H3 were used as loading control) CHD8 and H3K36me3 values relative to Sh-*GFP* controls. Mean values ± s.e. from independent biological replicates (n=4 for Sh4-*CHD8* and n=6 for the other samples) are plotted. t test for two mean population was performed. * p≤0.05 E. Bar plot represents GO biological process terms significantly enriched in genes with a reduced level of H3K36me3 at transcriptional elongation sites upon *CHD8* suppression. The bars are ordered according to adjusted p values in -log10 scale, the x-axis represents the number of genes enriched for each term. Top 20 GO terms are shown in figure, while full list of significant terms is reported in Suppl. Table 1.

Genes presenting reduced peaks for H3K4me1, H3K27ac and H3K36me3 histone marks were involved in “protein targeting”, “ribosome biogenesis” and “mitotic nuclear division” pathways (Fig. 1D) and enriched for ‘essential genes’ (required for a cell’s survival, Wang et al. 2015) and ‘constrained’ genes (intolerant to loss of function mutations, gnomAD), ‘FMRP targets in brain’ (Darnell et al. 2011), ‘post synaptic genes’ (Krishnan et al. 2016), and the ‘M3 co-expression module’ (Parikshak et al. 2013), whose expression peaks early during nervous system development (Fig. 1E).

### CHD8 suppression significantly affects transcriptional elongation chromatin states

By combined analysis of the histone marks enrichment through ChromHMM (Ernst and Kellis 2012), we defined 10 types of genomic regions with a specific chromatin state in control hiNPC (1. transcriptional initiation (H3K4me3, H3K4me2, H3K4me1, H3K36me3), 2. transcriptional elongation (high H3K36me3), 3. weak transcribed (low H3K36me3), 4. strong enhancer (H3K4me2, H3K4me1, H3K27ac), 5. weak/poised enhancer a (H3K4me2, H3K4me1), 6. weak/poised enhancer b (H3K4me1, H3K27ac), 7. active promoter (H3K4me3, H3K4me2, H3K4me1, H3K27ac), 8. inactive/poised promoter (H3K27me3, H3K4me3, H3K4me2, H3K4me1), 9. Polycomb repressed (H3K27me3), 10. heterochromatin/low signal (no enrichment)) (Fig. 2 A-left). To identify states affected by *CHD8* knock-down, we then compared the histone marks (peak counts for each histone mark within each chromatin states) across two conditions, i.e. controls versus *CHD8* knock-down (Sh2-*CHD8* and Sh4-*CHD8*) (Fig. 2 A-right) and detected transcriptional elongation, strong-weak/poised enhancer and active promoter as the chromatin states most affected by *CHD8* suppression.

To further dissect these results, we compared the number of peaks for each histone mark in controls and *CHD8* knock-downs within each chromatin state. While differences between biological replicates did not support a statistically significant variation at enhancer and promoter states (Supp. Fig. 3), the number of peaks decorated by histone H3K36me3 at transcriptional elongation genomic regions was significantly lower in Sh2-Sh4 *CHD8* (p value < 0.05, t test) compared to controls (Fig. 2B). Within the transcriptional elongation regions, H3K36me3 reduction affected ∼50% of all expressed genes in human neuronal progenitors (out of protein coding genes with > 2 TPM: 5447 lose H3K36me3, while 6451 were not affected). Additionally, genes that lost H3K36me3 following CHD8-haploinsufficiency, are significantly longer (90290 vs 60902 bp) and composed of larger exon number (7.36 vs 6.20) (not shown), compared to the unaffected ones. Importantly, independent western blot quantification of total histone H3K36me3 levels in the three Sh1-Sh2-Sh4 *CHD8* knock-down clones confirmed reduction of this histone modification compared to control Sh-*GFP* (Fig. 2 C-D). Genes with reduced H3K36me3 following *CHD8* suppression at transcriptional elongation sites were strongly enriched for processes related to cell cycle and proliferation (“cell cycle”, cell cycle process”, “mitotic cell cycle”, “ mitotic cell cycle process”) and “cell response to DNA damage stimulus” biological processes (Fig. 2E). For instance, the *HCN1*, gene encoding for a hyperpolarization gated chloride channel involved in epilepsy (Nava et al. 2014; Marini et al. 2018), presents reduced H3K36me3 enrichment following *CHD8* suppression, while other actively transcribed genomic locations (i.g. *BIRC6*) showed unaltered distribution (Suppl. Fig.4).

### CHD8 suppression-dependent reduction in Histone H3 Lysine 36 trimethylation impacts on CHD8-bound genes

By overlaying human CHD8 binding sites on the previously established chromatin states (Fig. 2A), we confirmed that CHD8 was confined to active promoters (90.11%) and, less prominently, to enhancers (strong, weak/poised a and b (3.18%, 3.00%, and 1.34%)) (Suppl. Fig. 5A) (Sugathan et al. 2014). As previously reported (Wade et al. 2018), CHD8 binding correlated with higher histone H3K36me3 (Suppl. Fig. 5B-E) and H3K4me3 enrichment metagene profiles (Suppl. Fig. 5C-F), as well as elevated RNA expression levels compared to CHD8-unbound genes (Suppl. Fig. 5 D).

Upon *CHD8* suppression, stringently defined (see materials and methods) CHD8-bound genes appeared to be more sensitive than CHD8-unbound genes (988 and 4205 respectively), presenting a significantly reduced H3K36me3 enrichment profile as confirmed by the effect size analysis (Fig. 3A-B. Suppl. Fig. 6A). Notably, the reduction in histone H3K36me3 elicited by *CHD8* suppression was specific since histone H3K4me3, another histone modification enriched at transcriptional start sites (TSS) of highly expressed genes (Suppl. Fig. 7A-B), remained unaltered (Suppl. Fig. 7C-E).

**Figure 3.**
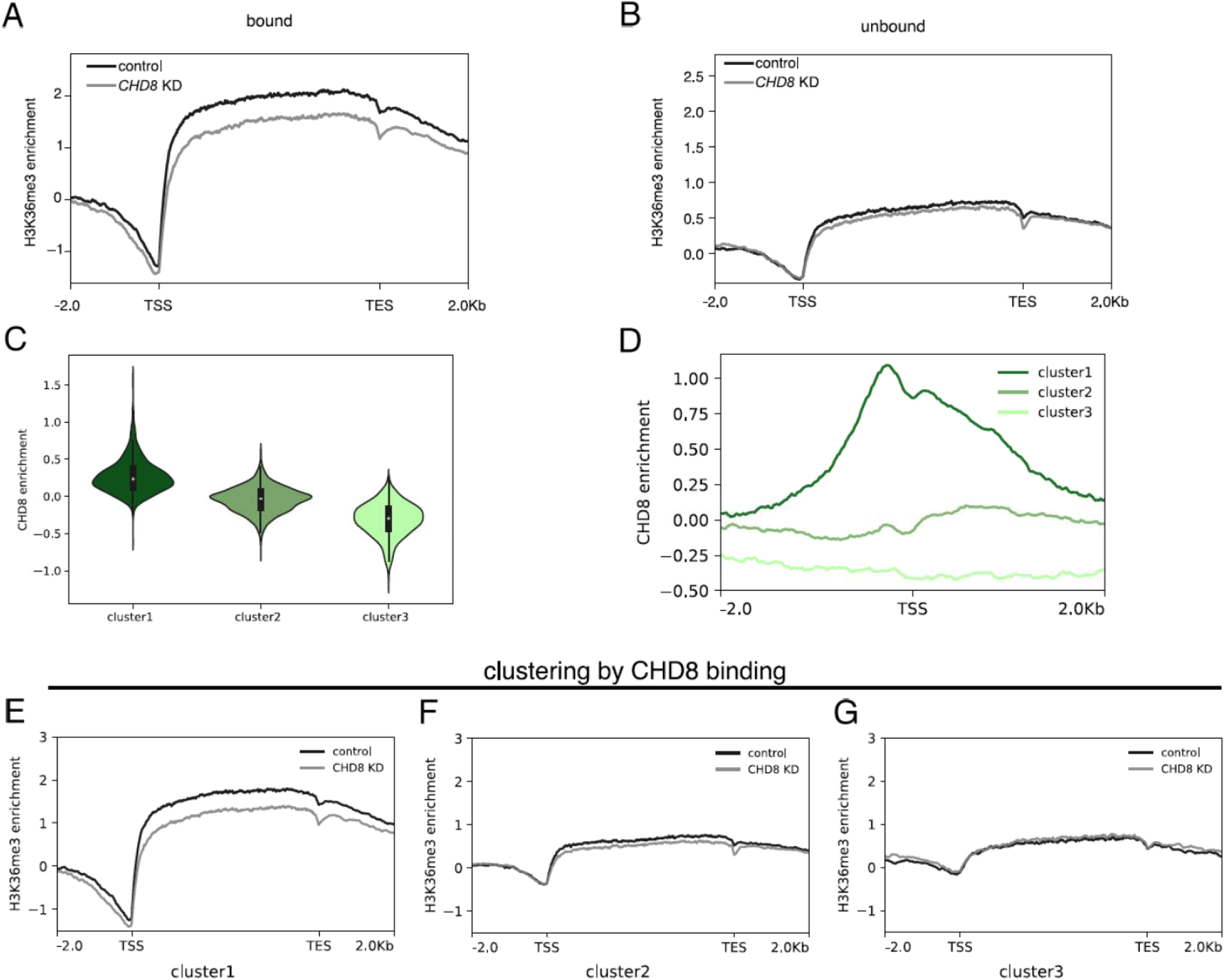
CHD8-suppression correlates with reduced H3K36me3 enrichment preferentially at CHD8-bound genes. A. B. Metagene profiles display the average of histone H3K36me3 enrichment (scaled log2 ratio of normalized ChIP value over INPUT control - see also Materials and Methods) in a region of ±2 Kbp upstream the transcriptional start site (TSS) and downstream the transcriptional end site (TES), calculated for control (black line) and *CHD8* knock-down (grey line) hiNPC and for CHD8-bound (#988) (A) and CHD8-unbound genes (#4205) (B). The difference between H3K36me3 enrichment in control and *CHD8* knock-down is significant for CHD8-bound genes, but not for CHD8 unbound genes (paired Cohen’s d effect size statistics). C. The violin plots represent the level of CHD8 binding enrichment (log2(ChIP/INPUT) for 3 groups of genes as clustered by k-means. Cluster #1 composed of 1239 genes shows high CHD8 enrichment (mean log2(ChIP/INPUT) = 0.27), cluster #2 composed of 2429 genes with medium-low CHD8 enrichment (mean log2(ChIP/INPUT) = −0.04) and cluster #3 composed of 1380 genes dispays negligible CHD8 enrichment (mean log2(ChIP/INPUT) = −0.31). D. Metagene profiles show the average of CHD8 binding enrichment in a region of ±2 Kbp around the TSS, calculated for the three clusters#1, #2, #3 as described in C. E. F. G. Metagene profiles display the average of histone H3K36me3 enrichment (log2(ChIP/INPUT)) in a region of ±2 Kbp upstream the TSS and downstream the TES, calculated for control (black line) and *CHD8* knock-down (grey line) hiNPC for each of the 3 clusters identified in Fig. 3C. The difference between H3K36me3 enrichment in control and *CHD8* knock-down is significant for Cluster#1, but not for Cluster#2 or Cluster#3 (paired Cohen’s d effect size statistics, see suppl. Fig. 6B-D).

Remarkably, clustering analysis based on CHD8 binding sites enrichment in control hiNPC identified 3 different clusters: Cluster #1 composed of 1239 genes with high CHD8 enrichment (mean log2(ChIP/INPUT) = 0.27), cluster #2 composed of 2429 genes with medium-low CHD8 enrichment (mean log2(ChIP/INPUT) = −0.04) and cluster #3 composed of 1380 genes with negligible CHD8 enrichment (mean log2(ChIP/INPUT) = −0.31) (Fig. 3C-D). Strikingly, clusters #1 was strongly affected by CHD8 decline, displaying significantly reduced H3K36me3 levels across the gene body (Fig. 3E and Suppl. Fig. 6B). Cluster #2 and #3, instead, with poor CHD8 enrichment in control hiNPC, displayed a correspondingly lower H3K36me3 enrichment, with no significant difference following CHD8 suppression (Fig. 3F-G and Suppl. Fig. 6C-D). In conclusion, this analysis confirmed that CHD8-bound genes were strongly sensitive to CHD8 reduction, presenting a substantial and specific drop in H3K36me3 histone modification (Fig. 3 C-E, Suppl. Fig.6C-D). Functional enrichment of genes from Clusters #1, i.e. CHD8-bound and losing H3K36me3 enrichment upon CHD8-suppression, highlighted GO Biological Process terms related to ‘mRNA processing’ and ‘RNA splicing’ (Suppl. Fig. 8 and Suppl. Table 1).

### Histone H3 Lysine 36 trimethylation elicited by CHD8 suppression alters RNA alternative splicing

To gauge the functional significance of H3K36me3 reduction observed following *CHD8* suppression, we then leveraged RNA-seq data from controls and *CHD8*-knockdown clones. As histone H3K36me3 seems to be correlated with high levels of RNA expression (suppl. Fig. 7A) (Wagner and Carpenter 2012), we reasoned that a decline in H3K36me3 levels would be associated with reduced RNA expression levels. Unexpectedly, however, haploinsufficient levels of CHD8 and, impaired H3K36me3 enrichment, didn’t correspond to a global difference in transcription in either CHD8-bound or CHD8-unbound genes (Fig. 4A-B).

**Figure 4.**
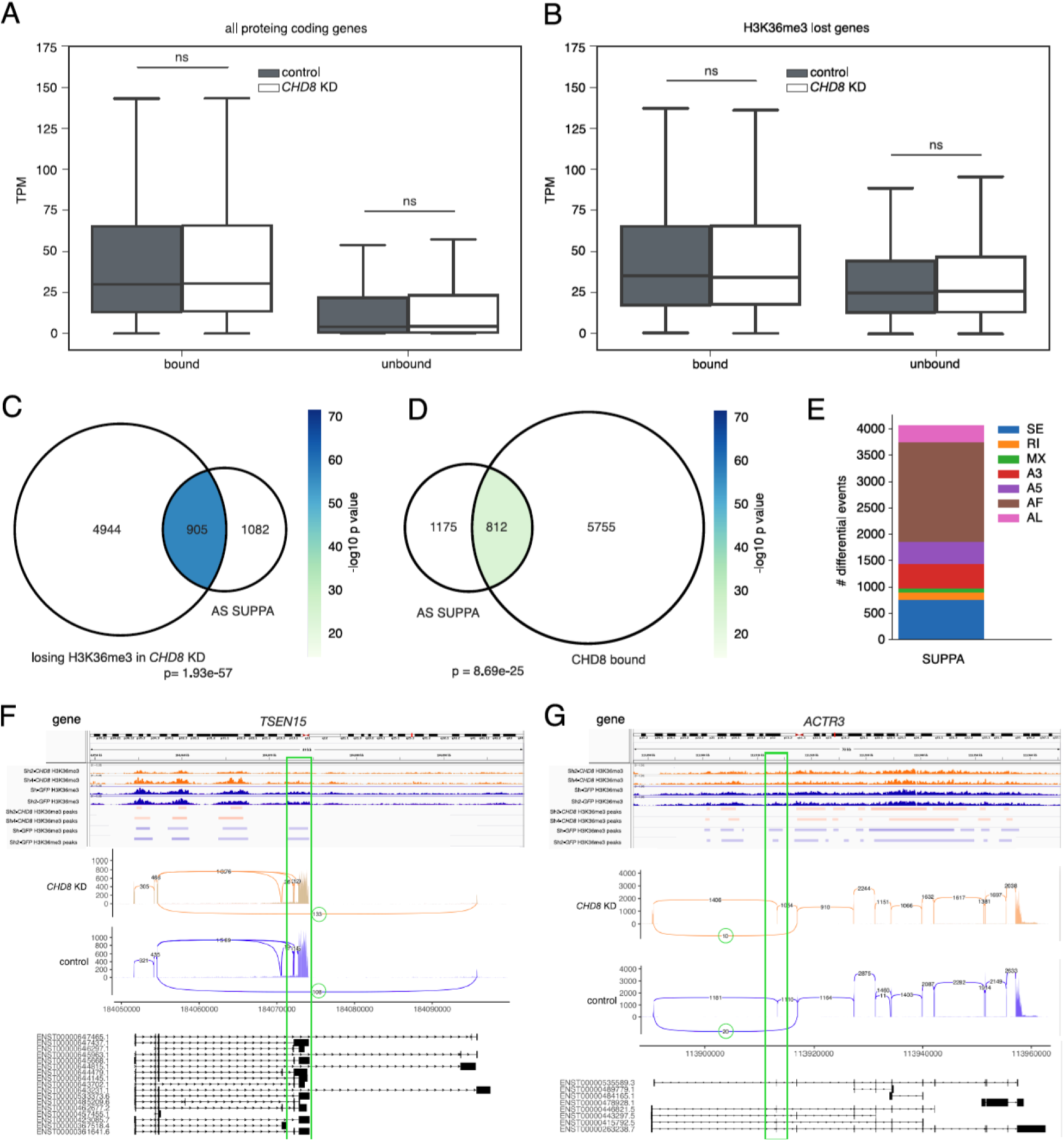
CHD8-suppression elicited reduction in H3K36me3 correlates with significant alterations in RNA alternative splicing. A. B. The box plots show the RNA expression level average of protein coding genes (A) and for genes that lose H3K36me3 peaks following *CHD8* knock-down (B). Control condition is shown in dark grey, *CHD8* knock-down in white; genes bound (left) and unbound by CHD8 (right) are also indicated. TPM: Transcripts Per Kilobase Million. Extreme outliers not shown. ns = not significant, t test statistic. C. D. Venn diagrams represent the overlap between genes losing H3K36me3 peaks following *CHD8* knock-down (losing H3K36me3 in *CHD8* knock-down) and genes presenting altered alternative splicing events as detected by SUPPA (AS SUPPA) (C), and the overlap between genes bound by CHD8 (CHD8-bound) and genes presenting altered alternative splicing events as detected by SUPPA (AS SUPPA). Number of genes for each condition is indicated. The enrichment significance for each intersection is computed by Fisher’s exact test and represented by colors. Color coded legend: -log10(p value). E. Stacked bar plot represents the 1987 differential alternative splicing events detected by SUPPA, distributed by event type. SE, skipped event; RI, retained intron; MX, mixed event; A3, alternative 3’; A5, alternative 5’; AF, alternative first exon; AL, alternative last exon. F. G. Data from two representative genes are shown along with their genomic sites, *TSEN15* (G) and *ACTR3* (F), including H3K36me3 ChIP-seq tracks for *CHD8* knock-down (Sh2-*CHD8* and Sh4-*CHD8* in orange) and control (Sh-*GFP* and Sh-*GFP2* in blue), with the corresponding peaks identified by MACS2 (top). At the bottom, sashimi plots display the alternative last (AL) event for *TSEN15* (G) and skipped exon (SE) for *ACTR3* (F), corresponding to the H3K36me3 peak lost after *CHD8* suppression (highlighted by green rectangles), in orange for *CHD8* knock-down and in blue for control.

Importantly, a significant proportion of genes losing H3K36me3 upon *CHD8*-suppression (905 genes of which 326 (36%) with CHD8 binding sites in promoter or enhancer regions) presented altered alternative splicing profiles, as evidenced by two analysis approaches (Fig. 4C-D and Suppl. Fig. 9) (Trincado et al. 2018; Shen et al. 2014). We noted an almost equal distribution of the H3K36me3 peaks lost following *CHD8* suppression between exonic and intronic regions (data not shown). While a significant correlation between reduction in H3K36me3 and alternative splicing was observed, however, the direct comparison of H3K36me3 lost peaks and differential splicing genomic coordinates revealed poor intersection. The vast majority of alternative splicing events with associated reduced H3K36me3 profiles were categorized as ‘alternative first exon’ (1892 (46.60%)) or ‘exon skipping’ (759 (18.69%)) (Fig. 4E,F and G). For the roughly ∼2000 differential splicing events detected by SUPPA (Fig. 4C,D), the proportion of genes presenting positive or negative ΔPSI (ΔPSI=PSI_ctrl - PSI_KD) (positive values: exon inclusion, negative values: exons skipping) remained pretty similar, however, for ‘exon skipping’ and ‘alternative 5’ events’ types (SE and A5), the majority (60%) PSI, presented a negative ΔPSI, suggesting that reduction of H3K36me3 could correlate with reduced exon inclusion/increased exon skipping. Among genes characterized by a reduction in H3K36me3 and presenting an altered splicing pattern following *CHD8* suppression, over-representation of GO terms and pathways related to “histone modification”, “covalent chromatin modification” and “mitotic cell cycle phase transition” (Suppl. Table 1) was observed.

## Discussion

Disruption of *CHD8* from *de novo* protein truncating variants and structural variants is well established as a highly penetrant risk factor for ASD (Neale et al. 2012; Satterstrom et al. 2020); O’Roak et al. 2012a; Talkowski et al. 2012). Chromodomain helicase DNA-binding protein 8 (CHD8), initially described as interacting with β-catenin as negative regulator of WNT signaling (Sakamoto et al. 2000; Nishiyama et al. 2004; Nishiyama et al. 2009; Nishiyama et al. 2012), has important roles during nervous system development. Mice completely lacking *Chd8* exhibit early embryonic lethality (between E5.5 and E7.5) (Nishiyama et al. 2009), while *Chd8* hypomorphic mutations are associated with perinatal mortality, pronounced brain hyperplasia in surviving animals and altered transcriptional expression (Hurley et al. 2018; Suetterlin et al. 2018). CHD8, a member of the chromo-helicases, has the ability to directly bind DNA (Sugathan et al. 2014; Cotney et al. 2015) and to slide nucleosomes (Manning and Yusufzai 2017). ChIP-sequencing has revealed ∼7000 CHD8 binding sites located at H3K27ac and H3K4me3 enriched regions, highly transcriptionally active promoters and enhancers (Sugathan et al. 2014; Cotney et al. 2015). However, with both direct and indirect transcriptional effects observed following *CHD8* suppression, its molecular mode of action in ASD remains unclear (Sugathan et al. 2014). CHD8 recruits histone KMT2/MLL methyltransferase complexes to induce mono-, di-, and trimethylation at lysine 4 of histone H3 and directly interacts with KMT2 components ASH2L and WDR5 (Thompson et al. 2008; Zhao et al. 2018). Reduced H3K27me3 enrichment was observed following *CHD8* suppression, which might reflect alteration to the core PRC2 methyltransferase, Ezh2 (Durak et al. 2016). However, a myriad of functions of CHD8 complicates this interpretation as it is associated with elongating RNAPII, so its role in transcriptional elongation has to be considered, especially for highly expressed genes, that are densely decorated by histone H3K36me3 (Rodriguez-Paredes et al. 2009).

In our ASD-relevant, human neuronal progenitor model system, we observed that *CHD8* suppression is prominently associated with a depletion rather than a gain, in H3K4me1, H3K27ac and H3K36me3 histone modifications. Genes displaying reduction in these histones marks are involved in ‘Ribosome Biogenesis’, ‘Mitotic cell Division’, ‘Methyltransferase Activity’ and strongly enriched for ‘Essential, Intolerant to LoF genes’, whose expression peaks early during nervous system development (M2, M3, post conception week 10-12) or increases during early cortical development (M16, post conception week 25-26) (Parikshak et al. 2013). The observed reduction in histone marks, while appreciable in all cases, is subtle and may suggest that CHD8 is unlikely to be a core component or a crucial recruiting factor for KMT2/COMPASS complex or SETD2/SETD5 methyltransferases. Rather it might act as a facilitator of histone methyltransferase activity. Using a combinatorial analysis of histone marks enrichment (Ernst and Kellis 2012), we dissected the differences elicited by *CHD8* suppression into 10 chromatin states. Stringent statistical criteria to focus on the strongest and most reliable epigenetic changes supported a drastic depletion of H3K36me3 peaks in the transcription elongation chromatin state. While an effect at enhancers and promoters could not be completely ruled out, CHD8 seems to facilitate the methyltransferase activity leading to H3K36 trimethylation. Given the dynamic balance between the activity of histone methyltransferases and demethylases, we also cannot exclude the hypothesis that CHD8 might work as an inhibitor of KDM2B (He et al. 2013). GO terms associated with ‘cell cycle progression’ and ‘mitosis’ are enriched among genes losing H3K36me3 at transcriptional elongation states. Interestingly, SETD2-5/KDM2B dysregulation has been correlated to the change of cell-cycle regulators (Kawakami et al. 2015), with altered H3K36 trimethylation associated with aberrant cell proliferation and neoplastic transformations (Wang et al. 2007; Jaffe et al. 2013; Simon et al. 2014). However, the modulation of H3K36 methylation is connected with the cell cycle not only in the context of tumorigenicity, but also during neuronal development and differentiation. Mutations in *SETD2* have been described in Sotos syndrome, childhood overgrowth condition with macrocephaly (Luscan et al. 2014) and in an ASD proband, also presenting macrocephaly (O’Roak et al. 2012a; Lumish et al. 2015), while disruptive mutations in *SETD5* – a newly described H3K36me3 methyltransferase (Sessa et al. 2019) - are associated with ID/ASD (Kuechler et al. 2015; Rauch et al., 2012; De Rubeis et al. 2014; Szczaluba et al. 2016; Parenti et al. 2017; Rawlins et al. 2017) and 3p25.3 microdeletion syndrome (Kellogg et al. 2013). Thus, it is possible that *CHD8* suppression, through its ability to modulate H3K36me3 levels, might lead to aberrant cell cycle regulation and macrocephaly as observed in animal models and in *CHD8*-autistic subjects (Bernier et al. 2014; Suetterlin et al. 2018). CHD8 direct targets, transcriptionally upregulated upon *CHD8* suppression (i.e. PML, HDAC7, CDK6 (Sugathan et al. 2014)) and implicated in cell cycle progression were indeed significantly enriched (Fisher’s exact test p value: 1.13×10^-7^) among genes that lose H3K36me3, suggesting a functional interplay between cell-cycle, CHD8 and H3K36me3.

But how can CHD8 modulate H3K36me3? From RNA sequencing data, *CHD8* suppression doesn’t seem to correlate with a direct reduction in SETD2/SETD5 levels nor to upregulation in H3K36me3 KDM2B methyltransferase (not shown). Based on our data, genes bound at their promoters/enhancers by CHD8 specifically present a significant depletion in H3K36me3, thus suggesting a possible direct interplay between the chromodomain and the H3K36 tri-methylases. Indeed, *SETD2* and *CHD8* display similar temporal expression patterns during human brain development (data from BrainSpan Atlas) (Bernier et al. 2014), though direct binding between CHD8 and SETD2 or SETD5 still needs to be demonstrated. The H3K36me3 depletion, observed upon *CHD8* suppression, is reminiscent of the function of another member of the CHD family, CHD1, which remodels nucleosomes within the gene body of actively transcribed gene (Petty and Pillus 2013). In fact, *chd1* loss alters H3K4me3 and H3K36me3 patterns throughout the yeast genome and loss of *chd1* causes increased cryptic transcription, altered splicing and nucleosome turnover within gene bodies (Petty and Pillus 2013; Lee et al. 2017). On the other hand, KIS-L in Drosophila is associated with all sites of transcriptionally active chromatin in a pattern that largely overlaps that of RNA Polymerase II (Pol II). Moreover, *kismet* mutant larvae, present severe reduction in the levels of elongating Pol II, suggesting that Kismet is also required for transcription elongation (Srinivasan et al. 2005). Similarly, it is tempting to hypothesize that CHD8, recruited to highly transcribed genes and in concert with SET2, might play crucial roles in nucleosome stability during elongation (Krogan et al. 2003) (Huang and Zhu 2018).

Trimethylation of H3K36 demarcates body regions of actively transcribed genes, providing signals for modulating transcription fidelity, mRNA splicing and DNA damage repair (Wagner and Carpenter 2012). Aberrant reduction in *SETD2* and reduced levels of H3K36me3 in our data are not directly causative of transcriptional differences and this is coherent with previous findings (Simon et al. 2014). Rather, reduced H3K36me3 correlates with altered alternative splicing. This is likely linked to the kinetics of PolII progression, since, H3K36me3 decorated nucleosomes act as intrinsic pause sites for elongating RNAPII. Altered H3K36me3 enrichment, then, could alter splice site recognition and change exon inclusion (Wagner and Carpenter 2012). In any event, alternative splicing differences detected in this study as a consequence of *CHD8* suppression, are particularly relevant in the context of nervous system development and function, especially for genes correlated with “RNA splicing” and “mRNA processing”. Interestingly, aberrant neuronal splicing has been previously related to *Chd8* haploinsufficiency in a mouse model of ASD (Gompers et al. 2017). In this context, our work establishing, for the first time, a functional link between H3K36me3 and *CHD8*, provides a molecular mechanism that warrants further investigations especially in neurodevelopmental syndromes and ASD in particular.

Finally, elongation-coupled H3K36 methylation usually serves also as a docking site for the histone deacetylase complex Rpd3S, which restores the repressive chromatin environment following Pol II passage to prevent cryptic transcription initiation (Huang and Zhu 2018). According to this hypothesis, *CHD8* suppression-related H3K36me3 reduction might also cause reduced repression in gene bodies, possibly resulting in increased uncontrolled, cryptic transcription. Although not possible to address in this study, which relied on previously generated poly-A plus RNA libraries, the analysis of spurious transcription as a consequence of H3K36me3 reduction, represents the next challenge for understanding chromatin-linked consequences in neurodevelopmental disorders and ASD in particular.

## Methods

### Cellular model

Human iPSC-derived NPC line GM8330-8 (Sheridan et al. 2011), Sh1-*CHD8*, Sh2-*CHD8*, Sh4-*CHD8* and Sh-*GFP* lines, previously generated by lentiviral delivery of shRNAs targeting *CHD8* and *GFP* coding sequences respectively (Sugathan et al. 2014), were kindly provided by Dr. Stephen Haggarty laboratory (Massachusetts General Hospital and Harvard Medical School, Boston, MA, U.S.A.).

Cells were cultured on poly-L-ornithine hydrobromide (20 µg/mL, Sigma)/laminin (3 µg/mL, Life Technologies) - coated plates in hiNPC medium (70% v/v DMEM (Life Technologies) completed with 30% v/v HAM F12 (Euroclone), 2% v/v B27 (Life Technologies), 1% v/v Penicillin-Streptomycin solution (Life Technologies) and 1% v/v L-Glutamine (Corning) and supplemented with EGF (20 ng/mL, Sigma), bFGF (20 ng/mL, R&D) and Heparin (5 µg/mL, Sigma)). Semi-confluent monolayers of hiNPCs were maintained in 5% CO2, 37°C humidified incubator.

### Protein extraction and western blot analysis

hiNPC cells were washed with PBS and resuspended in 100 ml of extraction buffer (10mM Hepes pH=8, 10mM KCl, 0.1mM MgCl2, 0.1mM EDTA pH=8, 0.1mM DTT and halt protease & phosphatase inhibitor cocktail (Life Technology)). Samples were centrifuged at 5.000 rpm for 10 minutes at 4°C to remove cytosolic fraction. Nuclear pellets were resuspended in HCl 0.2N and put in rotation at 4°C overnight. After centrifugation at 4.000 rpm for 10 minutes at 4°C, supernatants containing nuclear proteins were recovered. Proteins were quantified by Bradford Protein Assay Kit (Sigma-Aldrich). Protein samples were separated by 4-12% Bis-Tris Protein Gels (Thermo Fisher) and transferred on Amersham™ Protran™ 0.45µm Nitrocellulose (GE-healthcare) membrane. Membranes were blocked with 5% w/v nonfat dried milk and incubated with the following primary antibodies: anti-CHD8 (Novus Biologicals NB-10060417) (1:1000), anti-HSP90 (Cell Signaling Tech. #4874S) (1:5000), anti-Histone H3 (1:1.000) (Cell Signaling Tech, #4499), anti-Histone H3K36me3 (1:1.000) (Abcam #ab9050). Proteins were detected using horseradish peroxidase conjugated secondary antibodies anti-rabbit IgG 1:7.500 (GeneTex, #GTX213110-01), visualized by ECL Select WB detection reagent (GE Healthcare) following manufacturer’s instructions. Signal quantification was performed with Imagelab software (BIORAD).

### Statistical Analysis

The analysis of target proteins differences in western blotting experiments was evaluated by performing an unpaired, one-tailed t test. In all t tests, the significance level was set to 0.05. Data were represented as mean ± Standard Deviation (SD). The significance level was reported as: not significant (NS) p>0.05, * p≤0.05, ** p≤0.01, *** p≤0.001.

### Chromatin immunoprecipitation and sequencing (ChIP-seq)

Chromatin immunoprecipitation was performed using the protocol described by (Bernstein et al. 2006); with minor modifications. Briefly, ∼25 million iPSC-derived neural progenitor cells, controls and *CHD8*-Shs were fixed with 1% formaldehyde incubated for 10 min at RT with rotation. The crosslinking was quenched by adding 1.1 ml 2.5M Glycine, incubated 5 min at RT with rotation. The cells were pelleted at 1000 rpm and resuspended ice-cold PBS/protease inhibitor (PI) and spin for 5 minutes at 1000 rpm, washed with ice-cold PBS twice, harvested, pelleted and directly resuspended in 300ul lysis buffer/PI (50 mM Tris–HCl (pH8.1), 1% SDS, 10 mM EDTA), kept on ice for 10 minutes rotating occasionally and vortexed vigorously for 15 seconds every 3 minutes. Sonication of the samples 200-700 bps smear was accomplished using a Bioruptor sonicator (Diagenode), for a total of 45 minutes sonication at full power and sonication cycles of 30”ON/30”OFF. Samples were centrifuged at max speed for 10 minutes at 4°C. Then, sheared-chromatin was diluted 10 fold in ChIP dilution buffer (16.7 mM Tris–HCl (pH 8.1), 167 mM NaCl, 0.01% SDS, 1.1% Triton X-100, 1.2 mM EDTA), supplemented with protease inhibitor. 50 μl of sheared chromatin was removed and stored at 4°C as control aliquot (INPUT). Each sample was incubated at 4°C overnight with antibodies (20 ug/ChIP) of interest. The following primary antibodies were used: H3K27me3 (07-449, Millipore), H3K4me3 (AB8580, Abcam), H3K36me3 (Ab9050, Abcam), H3K4me2 (AB7766, Abcam), H3K4me1 (AB8895, Abcam) and H3K27ac (AB4729, Abcam). Chromatin–Antibody complexes were precipitated with Dynabeads Protein A beads (Invitrogen) and washed sequentially with low-salt (20 mM Tris–HCl (pH 8.1), 150 mM NaCl, 0.1% SDS, 1% Triton X-100, 2 mM EDTA), high-salt (20 mM Tris–HCl (pH 8.1), 500 mM NaCl, 0.1%SDS, 1% Triton X-100, 2 mM EDTA), LiCl (10 mM Tris–HCl (pH 8.1), 0.25 M LiCl, 1% NP40, 1% sodium deoxycholate,1 mM EDTA), and TE wash buffers (10 mM Tris– HCl (pH 8.0), 1 mM EDTA). Immunoprecipitated chromatin and INPUT samples were then eluted in elution buffer (TE plus 1% SDS, 150 mM NaCl, 5 mM DTT), de-crosslinked at 65°C for overnight and treated with proteinase K. DNA isolation was performed by phenol:chloroform:isoamyl alcohol. DNA was precipitated with 200 mM NaCl, supplemented with 30 μg glycogen, washed with EtOH and then treated with RNase I (Invitrogen). Finally, DNA was purified with MinElute Kit (Qiagen). Quantification of ChIP and INPUT DNA was accomplished using Qubit 2.0 Fluorometer system (Invitrogen). ChIP-seq libraries were prepared using NEBNext UltraII DNA Library preparation kit (Illumina) following manufacturer’s instruction with no modification. ChIP-seq libraries were prepared starting from 5ng of fragmented DNA using NEBNext UltraII DNA Library preparation kit (Illumina) following manufacturer’s instruction with no modifications. In order to obtain enough material for sequencing, eight cycles of PCR amplification were performed on adaptor ligated fragments.

### Histone marks ChIP-seq Data Analysis

ChIP-seq reads were aligned on the human genome reference assembly GRCh38 using BWA (version 0.7.15) (Li and Durbin 2009). Aligned reads were filtered to discard unmapped, multiply mapped, PCR duplicates reads (Picard tools MarkDuplicates version: 2.3.0, Picard Toolkit.” 2019. Broad Institute, GitHub Repository. http://broadinstitute.github.io/picard/) along with low quality alignments (samtools view -q 1, samtools version 1.7) (Li et al. 2009). Peak calling was performed with MACS2 (version 2.1.0) using a minimum FDR threshold 0.00001. The same settings with the addition of the --broad option were used for H3K27me3 and H3K36me3 marks (Zhang et al. 2008). Peaks localized to blacklisted regions (http://mitra.stanford.edu/kundaje/akundaje/release/blacklists/) or unplaced contigs were filtered. Narrow peaks (for H3K27ac, H3K4me1, H3K4me2, H3K4me3 histone marks) closer than 350 bp were merged into a single peak. Peaks were considered common between replicates if they overlapped by at least 50% of the length of the shortest peak, then extended coordinates were maintained and used in downstream analyses. CHD8 ChIP-seq reads from Sugathan et al. 2014 were realigned to the reference GRCh38 and peaks were recalled with the same procedure used for the narrow histone marks, with a default FDR threshold 0.05 as only peaks identified by all three antibodies were retained. GENCODE v.26 was used for peaks annotation (Frankish et al. 2019). Genes losing H3K4me1, H3K27ac and H3K36me3 peaks in *CHD8* knock-down were tested for enrichment of GO terms with the enricher function from clusterProfiler (version 3.10.1, Yu et al. 2012) with Benjamini-Hochberg adjusted p value cutoff of 0.05. The full list of slimGO terms used was downloaded from Ensembl BioMart on September 9^th^, 2019. Enrichment of the same gene list was assessed on custom gene sets by using one-tailed Fisher’s exact test. Custom gene sets were derived from public databases and publications of interest (Supplementary Table 1). Only non-redundant gene lists enriched at p value < 0.01 were shown in Fig. 1E, complete enrichment results are available in Supplementary Table 1. Combining the histone marks enrichment patterns over the genome, 10 chromatin states were identified for control (Sh-*GFP*, Sh-*GFP*2) using ChromHMM (version 1.14) (Ernst and Kellis 2012). The states were manually annotated according to the literature. The number of peaks called in these regions in both replicates for each histone mark was counted using BEDTools intersect, version 2.25.0 (Quinlan and Hall 2010), and the difference between control and *CHD8* knock-down was calculated. The total number of peaks for each histone marks in percentage was plotted as heat map in control cells (Fig. 2A), the difference (Fig. 2B) refers to the percentage of control peaks – *CHD8* KD peaks. The number of peaks per mark called in each chromatin state was tested with a two-sided t test considering two replicates in both control and *CHD8* knock-down. To generate the metagene profiles with deepTools, version 3.2.1, (Ramirez et al. 2016), ChIP-seq samples were normalized to INPUT with the SES method (Diaz et al. 2012). Enrichment was calculated in 10 bp bins over the gene body, scaled to 5 Kbp, and 2 Kbp up- and down-stream of the gene. In order to minimize noise in the metagene profile plots, a stringent set of filters was applied to protein coding genes before plotting: a minimum length of 2 Kbp, a minimum distance of 4 Kbp from other genes and absence of other features on the opposite strand, leading to a set of 9442 protein coding genes. For each histone mark, only genes with enrichment were plotted (at least one non-zero bin). These genes were divided into three groups based on *CHD8* binding enrichment pattern via k-means with deepTools plotProfile command. To complement the statistical hypothesis testing, paired Cohen’s d effect size statistics were calculated between groups along the entire region (Gibbons 1993). 99% simultaneous confidence intervals were constructed controlling FWER (Bonferroni correction). CHD8 binding sites profiles were calculated with deepTools computeMatrix reference-point and plotted with deepTools plotProfile. Visualization of enrichment tracks for chromatin was performed with the Integrative Genomic Viewer (IGV, version 2.4.9 (Robinson et al. 2011)).

### RNA-seq Data Analysis

Raw reads obtained from Sugathan et al., 2014 for corresponding samples (Sh-*GFP*, Sh-*GFP2*, Sh2-*CHD8*, Sh4-*CHD8*) were used to calculate transcripts abundance by kallisto (version 0.44.0) (Bray et al. 2016) on GENCODE v.26 transcripts. Transcripts per Million (TPM) were used to plot expression levels. SUPPA (version 2.3) was used to calculate the percent spliced-in (PSI) value per splicing event with an empirical method (Trincado et al. 2018). The same raw reads from Sugathan et al. 2014 were aligned with STAR (version 2.6) (Dobin et al. 2013) with default parameters on the GRCh38 reference, to analyze the alternative splicing with rMATS (version 3.1.0, Shen et al. 2014). As the two alternative splicing analysis methods are complementary (https://github.com/comprna/SUPPA/issues/47), both were included in the analysis. To check the overlap of splicing events with chromatin marks, the spliced in/out exon coordinates were intersected with the histone mark peaks’ coordinates. Alternative splicing events were represented in sashimi plots via ggsashimi (version 0.4.0) (Garrido-Martin et al. 2018) using GENCODE annotation v.33 as a reference (Frankish et al. 2019).

## Acknowledgments

We are indebted to the Biagioli, Ferrari and Talkowski lab members for helpful discussions, to Dr. Albert Basson (King’s College, UK) and Dr. Fulvio Chiacchiera (Department CIBIO, University of Trento) for critical reading of the manuscript. We are grateful to Dr. Alessandro Quattrone (Director, Department CIBIO, University of Trento) for the scientific support to CIBIO Core Facilities research. We thank Drs. Veronica De Sanctis and Roberto Bertorelli for technical support and assistance with preparation and sequencing of libraries. We thank Dr. Stephen Haggarty (Massachusetts General Hospital and Harvard Medical School, Boston, MA, U.S.A.) for sharing the human neuronal progenitor cell lines. This work was supported by Department CIBIO Institutional funding to M.B. and S.P, grants from Autism Speaks and the Simons Foundation Autism Research Initiative to J.F.G. and the National Institutes of Health GM061354 and NS093200 to J.F.G and M.E.T. E.S. was supported by the Structured International Post-doc Program of SEMM (SIPOD) and the AFM-TELETHON fellowship n. 21835.

## Author contributions

E.K, T.T., S.E., E.Salviato, F.D.L., E.S., M.A., performed experiments and analyzed the data. E.Salviato assisted with statistical analysis. E.S. and M.Benelli assisted with data analysis of alternative splicing and histone modifications. E.K. and M.Biagioli designed the study and wrote the paper. J.F.G., S.P., F.D., F.F., and M.E.T. provided intellectual guidance and assistance with bioinformatic analyses. M. Biagioli, M.E.T. and F.F. supervised this study. All authors discussed the results and commented on the manuscript.

