## Supplementary figures and images for "*CHD8* Suppression Impacts on Histone H3 Lysine 36 Trimethylation and Alters RNA Alternative Splicing"

### Supplemental Fig. 1

Kerschbamer_SupFig1

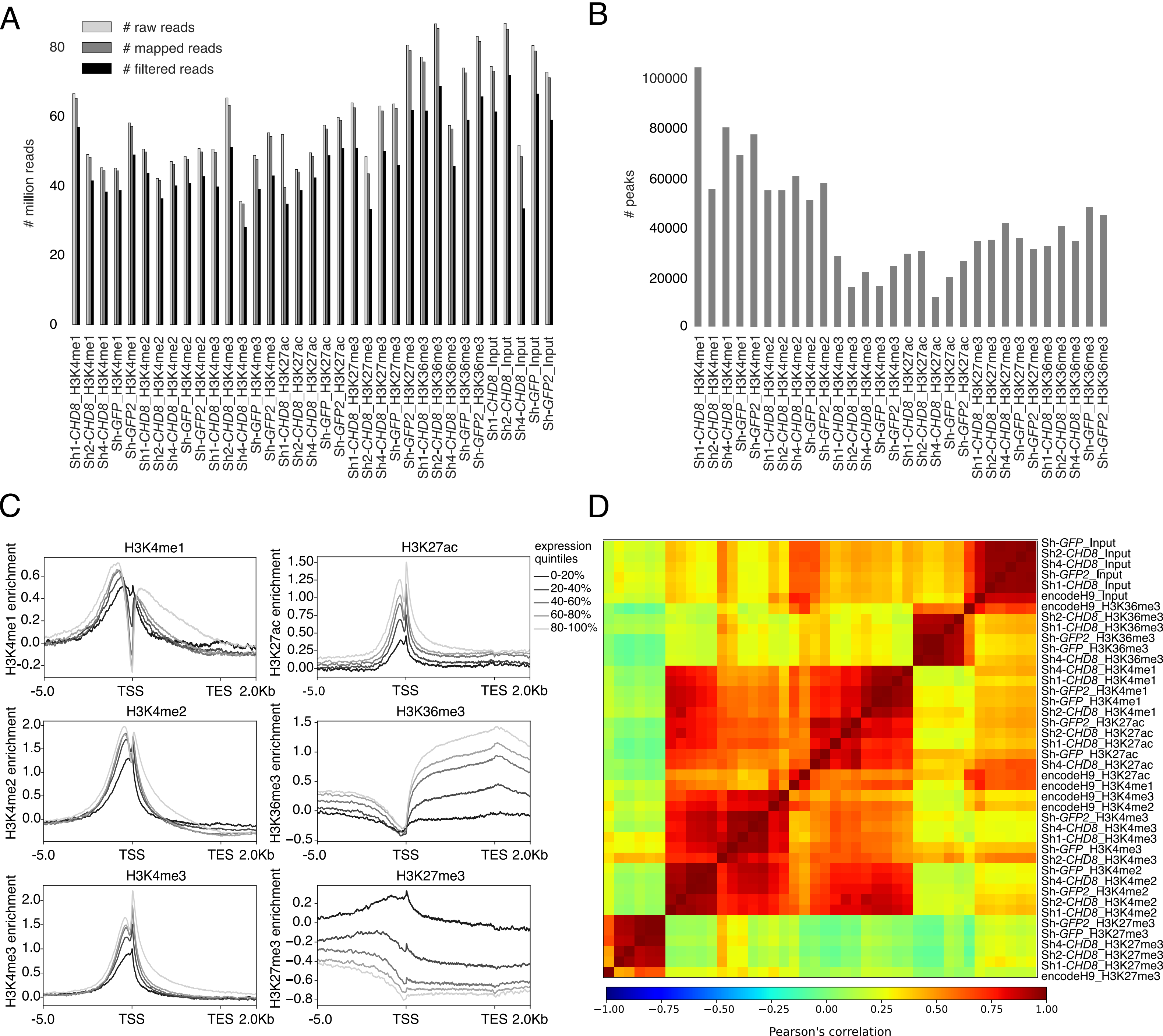

### Supplemental Fig. 2

Kerschbamer_SupFig2

A

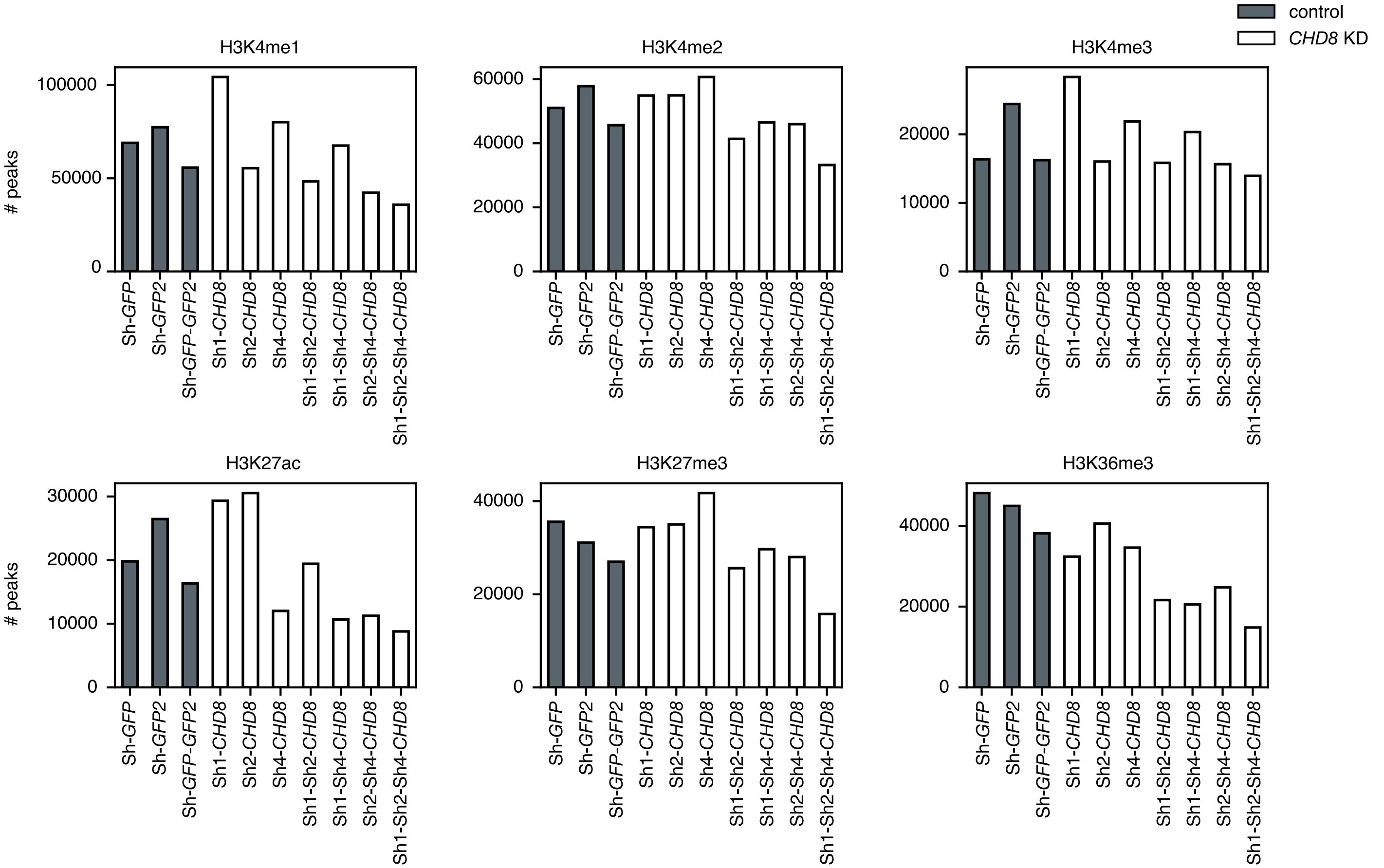


B

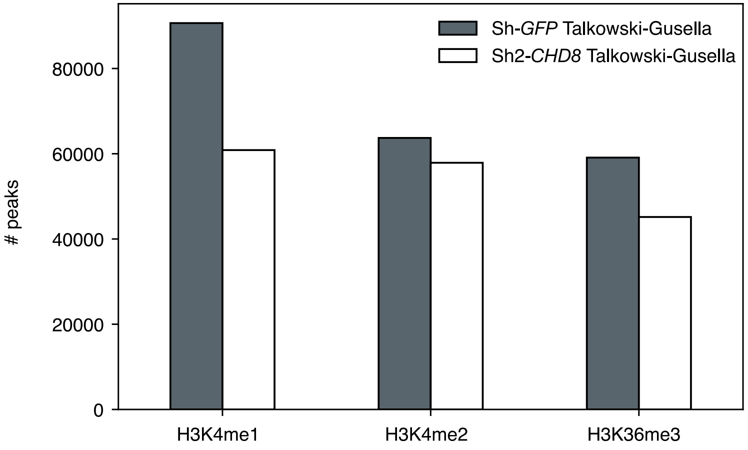

### Supplemental Fig. 3

Kerschbamer_SupFig3

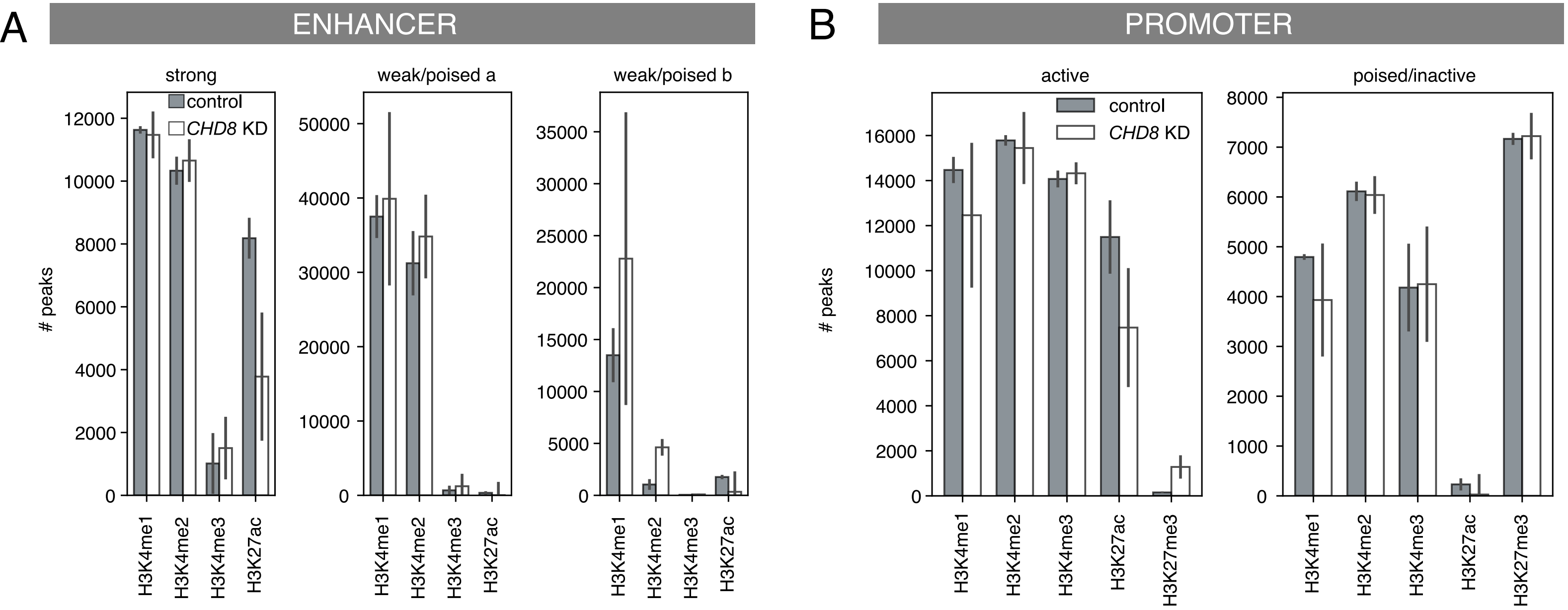

### Supplemental Fig. 4

Kerschbamer_SupFig4

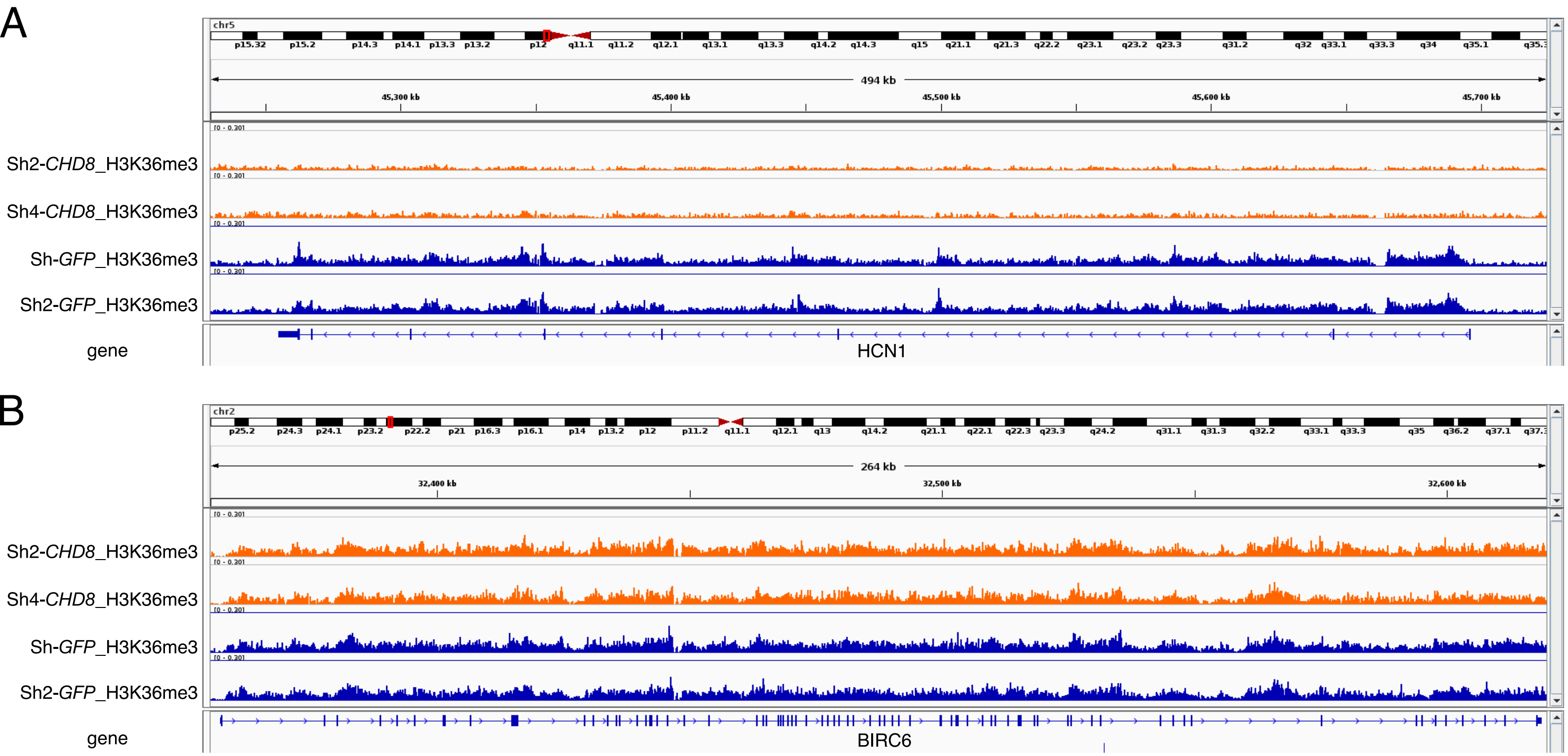

### Supplemental Fig. 5

Kerschbamer_SupFig5

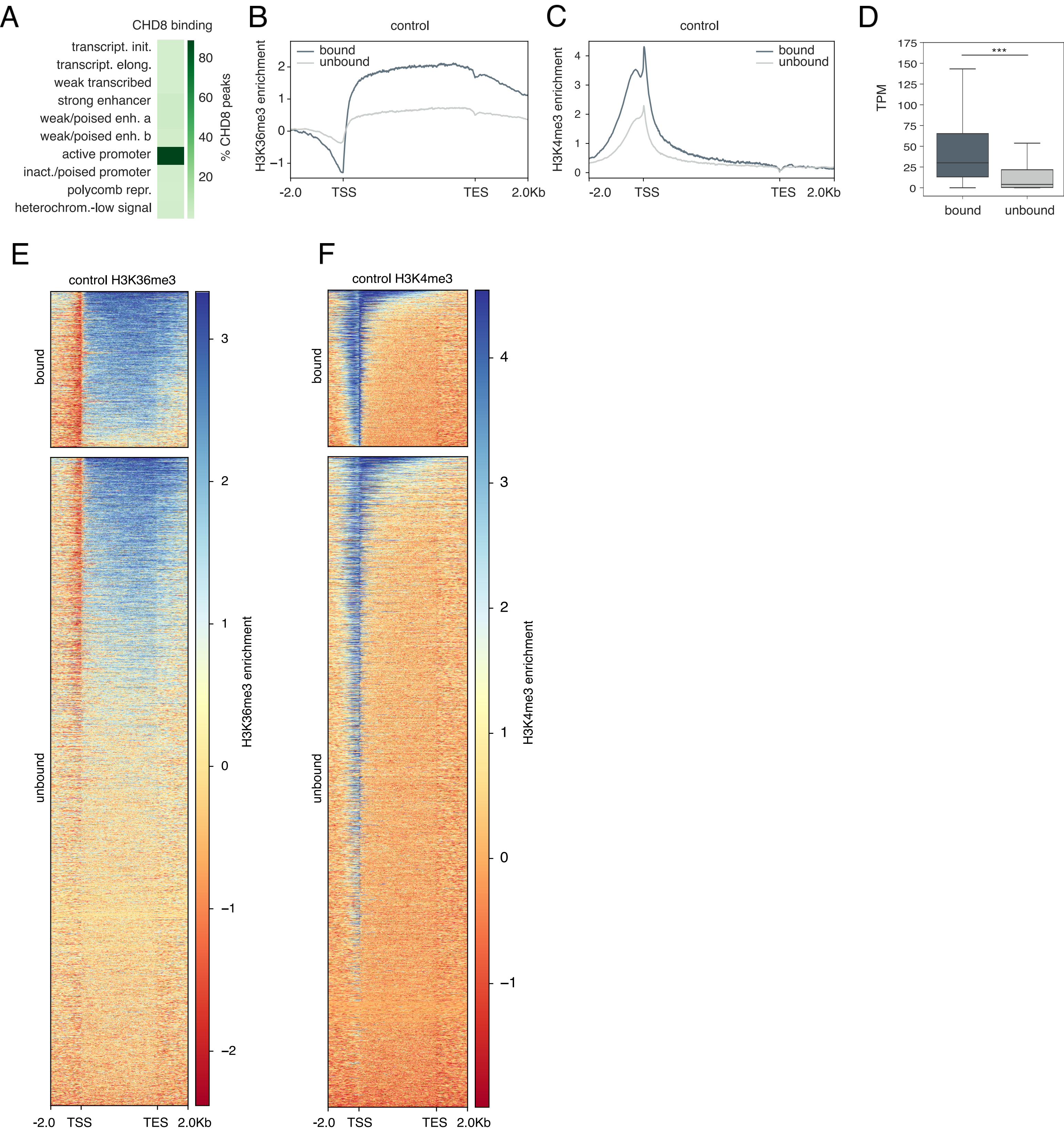

### Supplemental Fig. 6

Kerschbamer_SupFig6

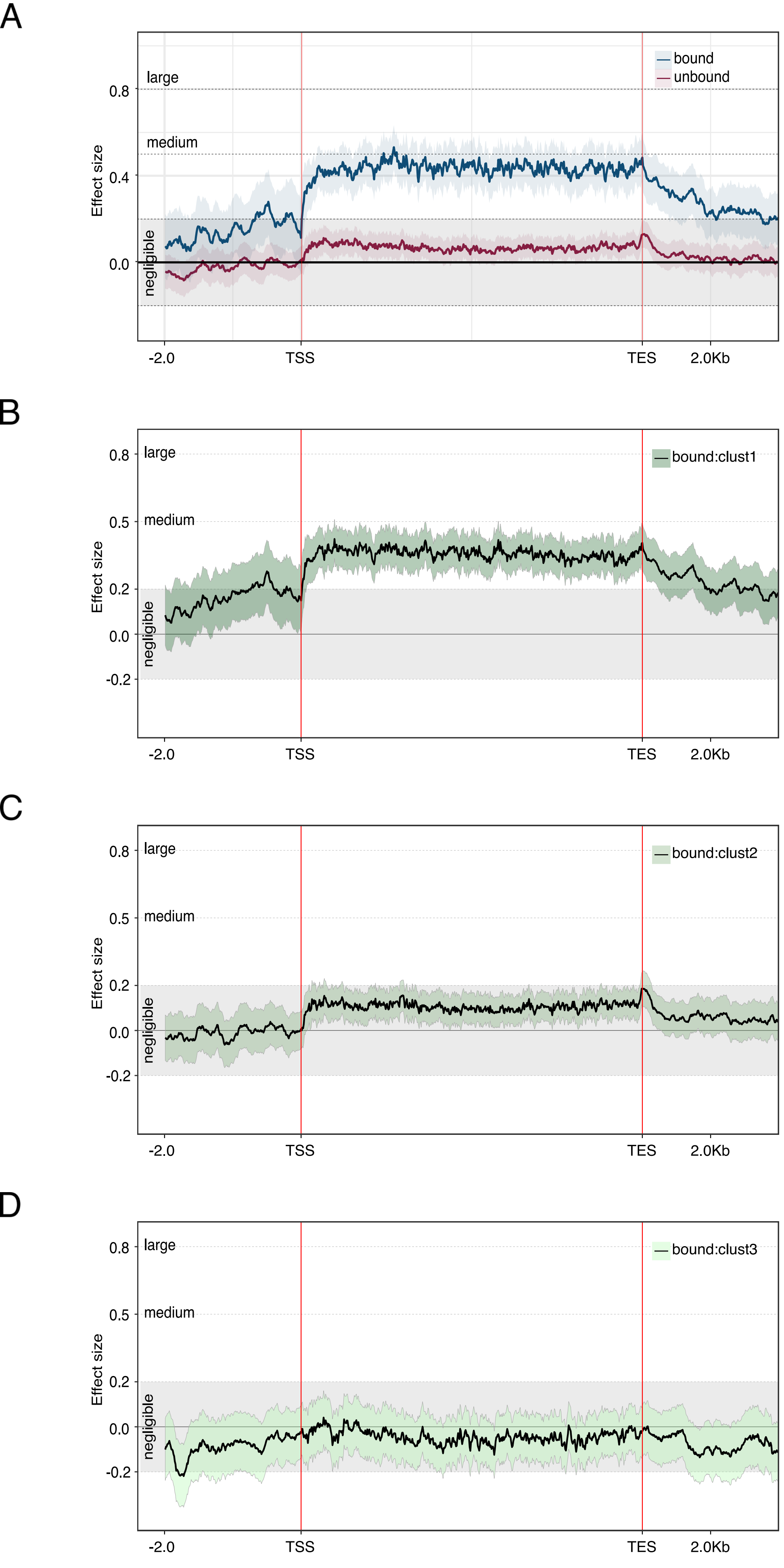

### Supplemental Fig. 7

Kerschbamer_SupFig7

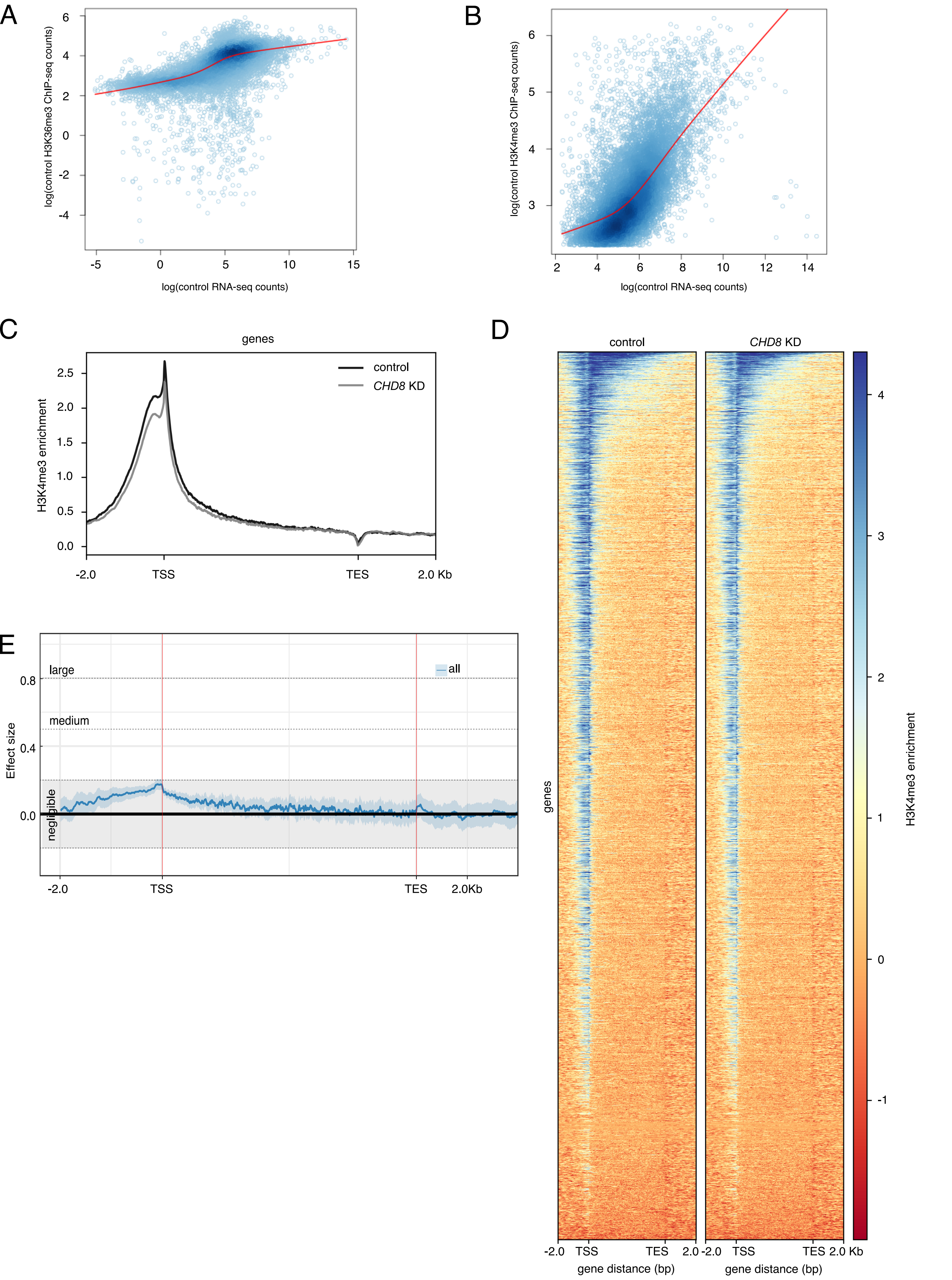

### Supplemental Fig. 8

#### Kerschbamer_SupFig8

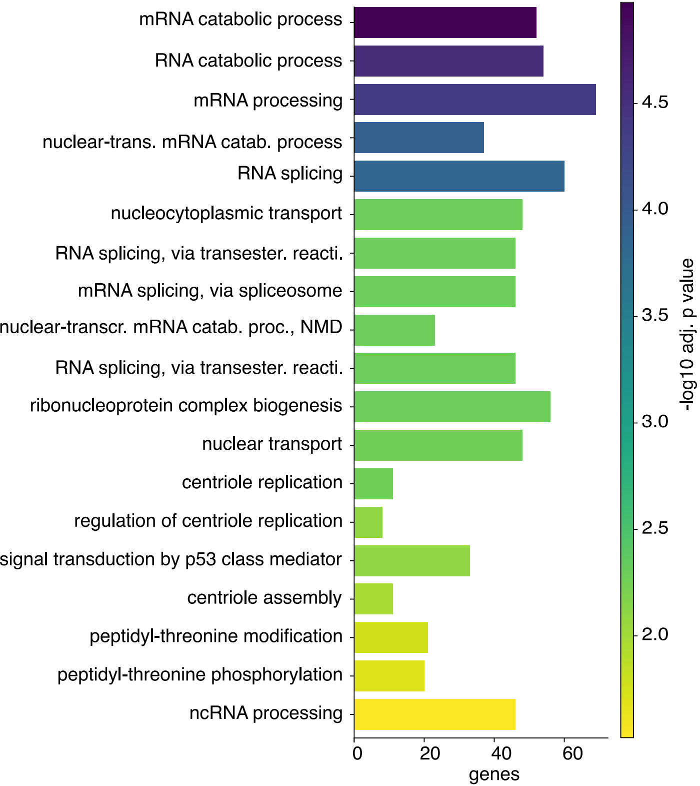

### Supplemental Fig. 9

Kerschbamer_SupFig9

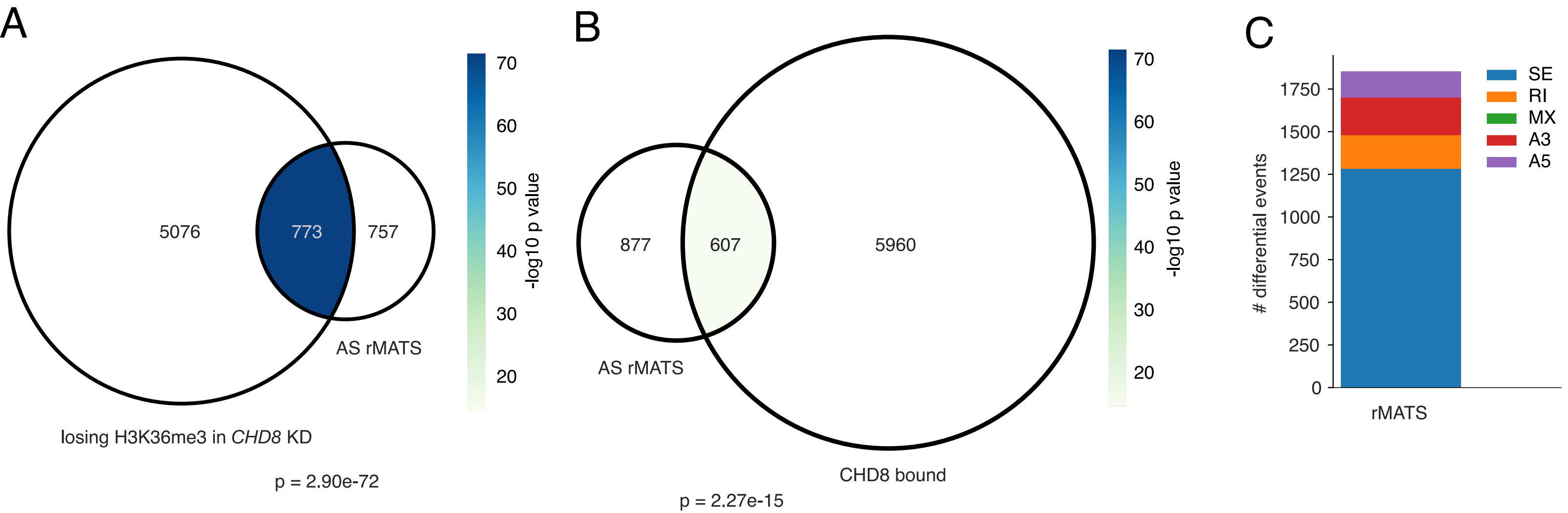
