## Supplemental Text for "*CHD8* Suppression Impacts on Histone H3 Lysine 36 Trimethylation and Alters RNA Alternative Splicing"

**Supplementary Methods**

Reads obtained from ChIP-seq libraries were checked for quality using FastQC (https://www.bioinformatics.babraham.ac.uk/projects/fastqc/) and MultiQC (<http://multiqc.info/>).

The metagene enrichment profile for each histone mark, was normalized against INPUT sample and plotted with deepTools grouping genes in five quintiles according to their level of expression as reported in Sugathan et al. 2014. ChIP-seq samples Spearman’s correlation on the enrichment pattern was calculated and plotted with deepTools, version 2.3.5, (Ramirez et al. 2016) on all our samples. As a control, the same histone marks ChIP-seq samples in neural progenitor cells (H9 derived) from ENCODE (Consortium 2012; Davis et al. 2018) were inserted in the correlation plot (ENCSR274OIJ, ENCSR645BCH, ENCSR661MUS, ENCSR449AXO, ENCSR139PIA, ENCSR573CWZ, ENCSR603SVD).

The same approach used for histone marks was carried on for H3K4me1, H3K4me2 and H3K36me3 Sh-*GFP* and Sh2-*CHD8* samples previously sequenced in the Talkowski-Gusella laboratories (Center for Genomic Medicine, Massachusetts General Hospital, Boston, MA, United States).

To check the correlation between histone modifications that mark actively transcribed regions and transcription data, (H3K4me3 and H3K36me3) ChIP-seq and RNA-seq reads were counted by BEDTools multicov, version 2.25.0, (Quinlan and Hall 2010). They were then normalized by library size and gene length, and displayed in a scatter plot. The LOWESS smoother was computed using R’s lowess function and plotted with R.

**Supplementary Figures**

***Supp. Figure 1 (related to Fig. 1)*** *Quality controls of read mappings and enrichment profiles.*

**A.** The bar plot represents the total number of raw (light grey), mapped (dark grey) and filtered reads (black) for each condition and histone mark.

**B.** The bar plot represents the total number of peaks called for each condition and histone mark. Details on peak calling were reported in Materials and Methods.

**C.** Metagene profiles of the six histone marks analyzed. Different outlines represent five groups of genes sorted based on their RNA expression levels. RNA expression from Sugathan et al. 2014 is calculated in quintiles and represented in shades of grey. Every histone mark displays the expected profile around the Transcription Start Site (TSS) or the gene body [TSS till Transcription End Site (TES)]. Highly expressed genes (80-100% expression quintiles, light grey line) show enrichment for histone marks associated with active transcription, while histone H3K27me3 shows a corresponding depletion.

**D.** The heatmap reports the Pearson’s correlation values calculated between different samples of this study and ENCODE datasets obtained from human neural progenitor cells [H9 cells ChIP-seq samples (details in Supplementary Methods)]. Correlation values are constructed on ChIP coverage calculated in 10 Kbp bins over the whole genome. Clustering is following the specific histone mark (multiple conditions and ENCODE datasets clusters together), confirming a similar enrichment pattern over the genome. Histone marks presenting enrichment at the transcription start site (H3K4me, H3K27ac) appear to be grouped together (central part of the heatmap).

***Supp. Figure 2*** ***(related to Fig. 1)*** *Peaks comparison across different CHD8 knock-down replicates and independent validation in ChIP-seq samples from Talkowski-Gusella laboratory (Massachusetts General Hospital, Boston, MA, United States.).*

**A.** The bar plots show the total number of peaks called in controls (Sh-*GFP*, Sh-*GFP2*, dark grey) and *CHD8* knock-down (Sh1-*CHD8*, Sh2-*CHD8*, Sh1-*CHD8,* white) for each of the histone marks analyzed. Sh-*GFP Sh-GFP2*, Sh1-Sh2 *CHD8*, Sh2-Sh4 *CHD8*, Sh1-Sh4 *CHD8* represent the intersection (number of peaks shared) of two biological replicates, while Sh1-Sh2-Sh4 *CHD8* indicates the intersection of three *CHD8* knock-down replicates. All intersections of two biological replicates - in any of the possible combinations - show the pattern described in Fig. 1, confirming a decreased number of peaks for H3K4me1, H3K27ac and H3K36me3 following *CHD8* suppression.

**B.** The bar plot describes the number of total peaks called in control (Sh-*GFP*, dark grey) and *CHD8* knock-down [Sh2-*CHD8* (Talkowski-Gusella), white] ChIP-seq samples for H3K4me1, H3K4me2, H3K36me3. These samples obtained and sequenced at a different time and in a different laboratory [*Talkowski-Gusella laboratory (Massachusetts General Hospital, Boston, MA, United States.)*], are confirming a specific decrease in histone H3K4me1 and H3K36me3 following *CHD8* down-regulation.

***Supp. Figure 3*** ***(related to Fig. 2)*** *CHD8 suppression does not significantly affect enhancer and promoter chromatin regions*

**A. B.** The bar graphs represent the number of peaks for each histone mark called at strong and weak/poised enhancers a/b genomic regions **(A)** and active and poised/inactive promoters **(B)**. Grey bars indicate controls (n=2, Sh-*GFP* and Sh-*GFP2*) and white bars refer to *CHD8* knock-down (n=2, Sh2-*CHD8* and Sh4-*CHD8*). Peaks differences between the two conditions at these chromatin states are not statistically significant (two-sided T test, see Materials and Methods for details).

***Supp. Figure 4 (related to Fig. 2)*** *Representative Integrative Genomic Viewer (IGV) snapshots of H3K36me3 enrichment at different genomic locations.*

**A. B.** Representative snapshots of the IGV (https://software.broadinstitute.org/software/igv/) genome browser views at the location of the hyperpolarization-activated, Epilepsy-associated cation channel *HCN1* (Nava et al. 2014; Marini et al. 2018) **(A)** and the cancer-associated, apoptosis-inhibitor *BIRC6* (Luk et al. 2017) **(B)** shows the ChIP-seq library-size normalized reads density (see Materials and Methods) for histone H3K36me3 across control (Sh-*GFP* and Sh-*GFP2*, blue tracks) and *CHD8* knock-down (Sh2-*CHD8* and Sh4-*CHD8*, orange tracks). At *HCN1* location, specific histone H3K36me3 depletion in *CHD8* knock-down samples compared to controls is observed, while at *BIRC6* locus, the expected enrichment for H3K36me3 histone mark is preserved in all conditions.

***Supp. Figure 5 (related to Fig. 3)*** *CHD8 binding correlates with high H3K36me3 and H3K4me3 and elevated RNA expression levels.*

**A.** The heatmaps represent 10 different chromatin states [1. transcriptional initiation, 2. transcriptional elongation, 3. weak transcribe, 4. strong enhancer, 5. weak/poised enhancer a, 6. weak/poised enhancer b, 7. active promoter, 8. inactive/poised promoter, 9. polycomb repressed, 10. heterochromatin/low signal] defined by the combination of different histone marks in control hiNPC as defined by ChromHMM(Ernst and Kellis 2012). The distribution of CHD8 binding sites (percentage of total CHD8 peaks) across different chromatin states (see Materials and Methods for details) is presented as percentage of the total and color-coded in the heatmap.

**B. C.** Metagene profiles display the average of histone H3K36me3 **(B)** and H3K4me3 **(C)** enrichment (scaled log2 ratio of normalized ChIP value over INPUT control - see also Materials and Methods) in a region of ±2 Kb around the gene body calculated for control hiNPC and for CHD8-bound (#988) and CHD8-unbound genes (#4205). The difference between histone H3K36me3 **(B)** and H3K4me3 **(C)** enrichment in CHD8-bound (dark grey) and unbound (light grey) genes is significant. TSS, Transcriptional Start Site; TES, Transcriptional End Site.

**D.** The box plot shows the average RNA expression level of protein coding genes bound (dark grey) and unbound by CHD8 (light grey). CHD8-bound genes correlate with higher expression levels. TPM: Transcripts Per Kilobase Million. Extreme outliers not displayed. *** = p <0.001, t test statistic.

**E. F.** Composite heatmaps plot the loci (rows) presenting H3K36me3 **(E)** and H3K4me3 enrichment **(F)** for CHD8-bound and CHD8-unbound genes. Genes in heatmaps are ranked based on their library-size normalized ChIP enrichment value relative to INPUT (enrichment score). Blue/red colors indicate high/low histone mark enrichment compared to INPUT.

***Supp. Figure 6 (related to Fig. 3)*** *CHD8-bound genes display a significantly different H3K36me3 enrichment along the gene body between control and* CHD8 *knock-down.*

**A.** Plot reporting Cohen’s d effect size statistics of the difference between control and *CHD8* knock-down H3K36me3 over 2 Kbp around the gene body of protein coding genes for CHD8-bound (blue) and CHD8-unbound genes (red) (transcriptional start and end sites are marked by red vertical lines). The effect size along the gene body is significant for CHD8-bound genes while it remains in the negligible area for CHD8-unbound genes.

**B.** Plot reporting Cohen’s d effect size statistics of the difference between control and *CHD8* knock-down H3K36me3 over 2 Kbp around the gene body of protein coding genes for Cluster#1 genes. Transcriptional start and end sites are marked by red vertical lines. The effect size along the gene body is significant.

**C. D.** Plot reporting Cohen’s d effect size statistics of the difference between control and *CHD8* knock-down H3K36me3 over 2 Kbp around the gene body of protein coding genes for Cluster#2 (**C.**) and Cluster#3 (**D.**). Transcriptional start and end sites are marked by red vertical lines. The effect size along the gene body is negligible for both clusters.

***Supp. Figure 7 (related to Fig. 3)*** *H3K4me3 and H3K36me3 correlation with RNA expression levels. Histone H3K4me3 enrichment is not affected by CHD8 knock-down.*

**A. B.** Scatter plots report the correlation between H3K36me3 ChIP-seq **(A)**, H3K4me3 ChIP-seq **(B)** (y-axis, normalized read counts) and RNA-seq (x-axis, normalized read counts) in control hiNPC. Lowess smooth is represented by red lines.

**C.** Metagene profile showing H3K4me3 enrichment (scaled log2 ratio of normalized ChIP/INPUT) in control (black line) and *CHD8* knock-down (grey line) on the TSS and along the gene body of protein coding genes (TSS, Transcriptional Start Site; TES, Transcriptional End Site).

**D.** Composite heatmaps plot the loci (rows) presenting H3K4me3 enrichment for protein coding genes in control (left) and *CHD8* knock-down (right). A region spanning ± 2Kb around the gene body is analyzed. Genes in heatmaps are ranked based on their library-size normalized ChIP enrichment value relative to INPUT (enrichment score). Blue/red colors indicate high/low histone H3K36me3 enrichment compared to INPUT.

**E.** Plot reporting Cohen’s d effect size statistics of the difference between control and *CHD8* knock-down H3K4me3 over 2 Kbp around the gene body of protein coding genes (transcriptional start and end sites are marked by red vertical lines). The effect size of the difference between H3K4me3 enrichment in control and *CHD8* knock-down along the whole region remains in the negligible area.

***Supp. Figure 8 (related to Fig. 3)*** *GO terms enrichment analysis for genes presenting lower H3K36me3 upon* CHD8 *suppression.*

Bar plot presenting the top 20 Biological Process GO terms significantly enriched in genes belonging to Cluster #, with high CHD8 binding enrichment in control hiNPCs and a lower H3K36me3 enrichment in *CHD8* knock-down (Fig. 3 E). Bars are colored according to -log10(adjusted p values) and x-axis represents the number of genes per term.

***Supp. Figure 9 (related to Fig. 4)*** *CHD8-suppression elicited reduction in H3K36me3 correlates with significant alterations in RNA alternative splicing – validation by rMATS, independent approach.*

**A. B.** Venn diagrams represent the overlap between genes losing H3K36me3 peaks following *CHD8* knock-down (losing H3K36me3 in *CHD8* KD) and genes presenting altered alternative splicing events as detected by rMATS (AS rMATS) **(A)**, and the overlap between genes bound by CHD8 (CHD8-bound) and genes presenting altered alternative splicing events as detected by rMATS (AS rMATS) **(B)**. Number of genes for each condition is indicated. The enrichment significance for each intersection is measured by Fisher’s exact test and represented by colors. Color coded legend: -log10(p value).

**C.** Stacked bar plot represents the 1484 differential alternative splicing events detected by rMATS, distributed by event type. SE, skipped event; RI, retained intron; MX, mixed event; A3, alternative 3’; A5, alternative 5’.
